# No evidence that individual alpha frequency (IAF) represents a mechanism underlying motion-position illusions

**DOI:** 10.1101/2024.07.17.603862

**Authors:** Timothy Cottier, William Turner, Violet J. Chae, Alex O. Holcombe, Hinze Hogendoorn

## Abstract

Motion-Position Illusions (MPIs) involve the position of an object being misperceived in the context of motion (i.e. when the object contains motion, is surrounded by motion, or is moving). A popular MPI is the flash-lag effect, where a static object briefly presented in spatiotemporal alignment with a moving object, is perceived in a position behind the moving object. Recently, Cottier et al. (2023) observed that there are stable individual differences in the magnitude of these illusions, and possibly even their direction. To investigate the possible neural correlates of these individual differences, the present study explored whether a trait-like component of brain activity, individual alpha frequency (IAF), could predict individual illusion magnitude. Previous reports have found some correlations between IAF and perceptual tasks. Participants (*N*=61) viewed the flash-lag effect (motion and luminance), Fröhlich effect, flash-drag effect, flash-grab effect, motion-induced position shift, twinkle-goes effect, and the flash-jump effect. In a separate session, five minutes of eyes-open and eyes-closed resting state EEG data was recorded. Correlation analyses revealed no evidence for a correlation between IAF and the magnitude of any MPIs. Overall, these results suggest that IAF does not represent a mechanism underlying MPIs, and that no single shared mechanism underlies these effects. This suggests that discrete sampling at alpha frequency is unlikely to be responsible for any of these illusions.

---

Motion-Position Illusions (MPIs) are a group of visual illusions in which the position of an object in the context of motion is incorrectly perceived. Typically, the object will contain internal motion, be surrounded by global motion, or the object itself will be in motion. The mechanisms underlying these illusions are highly debated and limited neural correlates have yet been identified. Recently, several studies have observed the presence of individual differences in the perception of MPIs (Cottier et al., 2023; Gauch & Kerzel, 2008; Morrow & Samaha, 2022). For some of these illusions, there is evidence that some participants consistently experience no illusory effect or the opposite of the expected effect. Individual differences often reflect differences in the optical and neural processes that mediate perception (Mollon et al., 2017). Therefore, by using an individual differences approach, we can elucidate the mechanisms contributing to these illusions and visual perception in general. This research is fundamentally important for understanding the basis of individual differences in motion and position perception.

As our perception of the world appears continuous, visual perception is typically assumed to be a continuous process. However, several researchers have argued that visual perception might in fact be discrete (Herzog et al., 2020; Menétrey et al., 2022; VanRullen, 2016; VanRullen & Koch, 2003; White, 2018). Similar to theories of discrete perception, discrete sampling is based upon the idea that visual input is sampled into discrete moments, and perception results from a reconstruction of several discrete perceptual moments (Schneider, 2018; Stroud, 1967). Schneider (2018) proposed a model of discrete sampling to explain various properties of the flash-lag effect, Fröhlich effect and related illusions.

The flash-lag effect (Figure 1A) involves briefly presenting a static object (the flash) in spatiotemporal alignment with a moving object (Nijhawan, 1994). While the two objects are physically aligned in time and space, the moving object is perceived in a position further along its motion trajectory, and the flashed object is perceived to lag behind. According to Schneider (2018) the flash-lag effect occurs because a moving object continues to move throughout a perceptual moment and is perceived as its last position in a given moment. Conversely, on average, the flash will have occurred prior to the end of the moment. When the flash is experienced at the end of the moment in its veridical position, the moving object will have progressed further along its trajectory, and will thus be experienced at a more advanced position. Schneider (2018) proposed that this discrete sampling and reconstruction process could correspond to alpha oscillations. However, this has yet to be tested.

**Figure 1.**
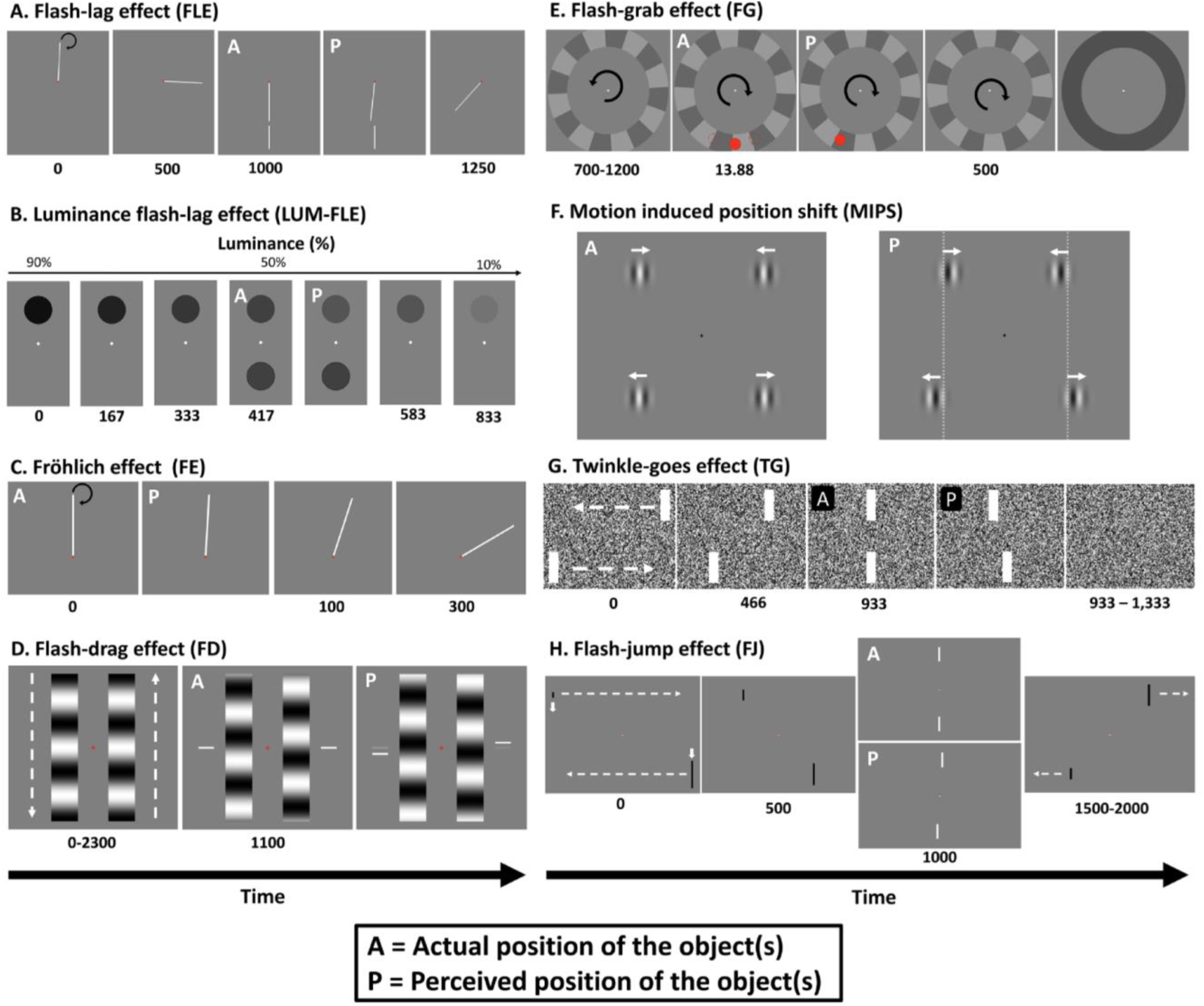
Image and caption reproduced with permission from Cottier et al. (2023, p.2), and consistent with their creative commons license. “Stylized depictions of example trials for the eight MPIs used in this study. Video examples for each illusion can be accessed at https://tcottier96.github.io. For all images, panels marked as “A” indicate the actual position of the object, and “P” indicates the perceived position of the object. (**A**) Flash-lag effect (FLE): a rod rotates clockwise around the fixation point for 1,250 ms. After 1 second, a stationary rod is briefly flashed in spatiotemporal alignment with the moving rod (actual). However, the moving rod is perceived mislocalized along its clockwise trajectory (perceived). (**B**) Luminance flash-lag effect (LUM-FLE): the top circle decreases in luminance over 833 ms. Halfway through the trial, on the opposite side of the fixation point, a circle with identical instantaneous luminance is briefly presented (actual). Even though both circles have identical luminance values, the target circle is perceived further along its luminance trajectory and thus is perceived to be brighter than the flashed circle (perceived). (**C**) Fröhlich effect (FE): a rod rotates clockwise around the fixation point. When the rod initially appears, it is pointing straight up (actual), but it will be perceived in a position along its clockwise trajectory (perceived). (**D**) Flash-drag (FD) effect: two sinusoidal gratings move in opposite directions for 2,300 ms. In this trial, the right grating is moving upward, while the left grating moves downward. After 1,100 ms, two bars are flashed on the outside of each grating. While these bars are presented in vertical alignment (actual), they are perceived mislocalized in the direction of their nearest grating’s motion (perceived). (**E**) Flash-grab effect (FG): an annulus rotates counterclockwise for 800 ms, then reverses direction and rotates counterclockwise for 500 ms before turning gray. At the moment the annulus reverses direction, a red circle is flashed for 13.88 ms in one of three positions (the dotted red lines). After the annulus turns gray, participants report the perceived location of the target with a mouse click. In this trial, the red circle was presented at the bottom center of the annulus (actual). However, this circle is perceived to be displaced in the reversal’s direction of motion (perceived). (**F**) Motion-induced position shift (MIPS): two pairs of vertically aligned gratings are presented (actual). The phase of the top gratings drifts toward the fixation point, while the phase of the bottom gratings drifts away from the fixation point. Even though the gratings are vertically aligned, they are perceived offset in their direction of motion (perceived). (**G**) Twinkle-goes effect (TG): two bars translate toward one another for 933 ms. The top bar is moving right to left, and the bottom bar is moving left to right. When the bars are vertically aligned (actual), they disappear on a background of dynamic noise. The perceived offset positions of the two bars are shifted forward along their respective trajectories, such that they are seen as misaligned (perceived). (**H**) Flash-jump effect (FJ): involves two bars moving toward each other and changing in height. In this trial, the top bar was moving right to left and increasing in height, while the bottom bar moved left to right while decreasing in height. When the two bars reach the center of the screen and are physically aligned, they will be the same height and briefly become white (actual). This brief color change is mislocalized further along the motion and growth trajectory of the bar and as such is perceived when the bar is a different size and not vertically aligned with the other bar (perceived).”

Alpha oscillations (7-13 Hz) are one of the most prominent brain rhythms in human neural recordings (Klimesch, 1999). Alpha oscillations predominantly occur over the occipital cortex, and thus are likely to reflect the sensory aspects of visual perception (VanRullen, 2016). Two studies have demonstrated how various features of alpha oscillations may influence the flash-lag effect, the most well-known MPI (Chakravarthi & VanRullen, 2012; Chota & VanRullen, 2019). In 2012, Chakravathi and VanRullen found a strong correlation between the flash-lag effect and pre-stimulus occipital theta and alpha phase between 5-10 Hz (with a peak at 7 Hz), and between high-alpha to low-beta band post-stimulus phase in the frontocentral electrodes (12-20 Hz). Consistent with these findings, Chota and VanRullen (2019) found that the flash-lag effect magnitude could be modulated by an entrainer oscillating at 10 Hz. These studies suggest that periodic alpha oscillations may modulate the perception of at least one MPI and thereby lend some support for the theory that discrete sampling underlies the flash-lag effect. However, these studies do not provide any insight into the extent to which alpha oscillations modulate perception of the broader class of MPIs, including the Fröhlich effect and flash-jump effect, which the perceptual sampling account attempts to account for (Schneider, 2018).

If alpha oscillations contribute to the perception of MPIs then alpha might predict individual differences in those illusions. Individual alpha frequency (IAF) is a trait-like component of alpha, with high heritability (Smit et al., 2006), that is unique to each individual and stable over time with excellent test-rest reliability (Grandy et al., 2013). IAF has been shown to correlate with general cognitive performance (Grandy et al., 2013), feature binding (Zhang et al., 2019) and spatial localisation (Howard et al., 2017).

Several studies have argued that IAF may index the temporal resolution of visual perception (for a review, see Samaha & Romei, 2024). Samaha and Postle (2015) found that IAF is related to whether two flashes presented in close proximity are perceived separately or instead fuse and are perceived as a single flash. They found that participants with a faster IAF could perceive both flashes at a shorter interstimulus interval than those with a slower IAF. On this basis, they argued that IAF is related to the segregation and integration of incoming sensory information, with individuals with faster IAF more able to segregate the two flashes as distinct entities at shorter interstimulus intervals. These influential findings have been replicated by several researchers (for a review, see Samaha & Romei, 2024), most recently by Deodato and Melcher (2024). These past studies thus provide solid evidence that IAF can reliably index individual differences in visual perception.

Empirical evidence has also emerged showing that IAF is related to the perception of illusions and motion. For example, IAF has been linked to perception of the sound-induced double flash illusion (Cecere et al., 2015), the bistable stream-bounce display (Ronconi et al., 2023), the perceived frequency of the illusory jitter in the motion-induced spatial conflict (Minami & Amano, 2017), the flickering wheel illusion (Sokoliuk & VanRullen, 2013), the spatial localisation of moving objects (Howard et al., 2017), and contrast detection abilities (Tarasi & Romei, 2024). Overall, these studies also suggest that IAF is related to individual differences in visual perception.

Regarding the flash-lag effect and Fröhlich effects in particular, Morrow and Samaha (2022) argued that if discrete sampling at alpha was contributing to the flash-lag and Fröhlich effects, then the illusion magnitudes of these effects should correlate with one another. This is based upon Schneider’s (2018) model, if one accepts that IAF indexes the duration of an individual’s perceptual moment. However, Morrow and Samaha (2022) did not find a correlation between the Fröhlich and flash-lag effects (*<u>r</u>*<u>_s_</u> = –.008, 95% CI = [-0.41, 0.39]), suggesting that these illusions are not caused by a shared underlying process. This finding could be a false negative, as their small sample size did not provide sufficient statistical power to detect weak-moderate effects. However, Cottier et al. (2023) also found that the correlation between the Fröhlich and flash-lag effects was close to zero (*r*_s_ = .1, 95% BCa CI = [-0.144, 0.336]), despite high individual task reliability and a much larger sample size. However, neither study analysed EEG to measure participants’ IAF and explore whether it correlated with individual illusions. Overall, the empirical evidence suggests that IAF is related to individual differences in visual perception, and aspects of alpha oscillations are related to the perception of the flash-lag effect (Chakravarthi & VanRullen, 2012; Chota & VanRullen, 2019). On this basis, we propose that IAF might be correlated with the perception of MPIs.

The present study assessed whether IAF is related to the magnitude of eight MPIs. This study was an extension of Cottier et al. (2023). As such, we adopted an individual differences approach and had participants complete the flash-lag effect (Nijhawan, 1994), luminance flash-lag effect (Sheth et al., 2000), Fröhlich effect (Fröhlich, 1924), flash-drag effect (Whitney & Cavanagh, 2000), flash-grab effect (Cavanagh & Anstis, 2013), motion-induced position shift (De Valois & De Valois, 1991), twinkle-goes effect (Nakayama & Holcombe, 2021), and flash-jump effect (Cai & Schlag, 2001). In order to calculate IAF, we also collected eyes-open and eyes-closed resting state EEG data in a separate experimental session. To briefly foreshadow our results, we find no evidence for a relationship between IAF and any of these illusions. This suggests that discrete sampling in the alpha range is unlikely to be responsible for MPIs. We also show that while we do not replicate the statistically significant correlations observed in Cottier et al. (2023) after correcting for multiple comparisons, our correlation estimates are nevertheless similar. As such, we conduct an auxiliary analysis which provides updated estimates of the inter-illusion correlation matrix, by pooling the data from Cottier et al. (2023) and the present study.

## Methods

### Participants

Cottier et al. (2023) found statistically significant correlations between certain MPIs of at least .37. Based on that, we used a Correlation: Bivariate normal model from the Exact test family (one-tailed) in G*power (version 3.1; Faul et al., 2009), to estimate a-priori that we required a sample size of 59 participants to have 90% power to detect such effects (alpha level = .05). Therefore, 61 participants aged between 18-51 (*M* = 25.6, *SD* = 6.89; 44 females) were recruited from the University of Melbourne’s paid research pool. Of these participants, 18 participated both in Cottier et al. (2023) and in a separate EEG study that recorded their resting state EEG. Participants were reimbursed $10/hr for the behavioural component of the study, and $15/hr for the EEG component. All participants self-reported as having correct or corrected to normal vision and no neurological deficits or disorders. Four participants reported being primarily left-handed, the remaining participants were right-handed. Some participants were excluded from analysis, which is discussed in detail in the pre-processing section below. This study was approved by the University of Melbourne’s Human Research Ethics committee, with separate approval provided for the illusion and EEG components (Illusion ID: 2022-12816-29275-8, EEG ID: 2022-12985-29276-6). Written informed consent was collected prior to participation.

### Apparatus

#### Behavioural experiment

Consistent with Cottier et al. (2023), stimuli were generated using PsychoPy (v2021.2.3; Peirce et al., 2019) and displayed upon a 24.5 ASUS PG258Q with a resolution of 1920 x 1080 pixels and a refresh rate of 144Hz. The experiment ran off an HP EliteDesk 800 G3 TWR Desktop PC with an Nvidia GTX 1060 graphics card, with the Windows operating system. The monitor was gamma corrected using a Cambridge Research Systems ColorCal MKII (Cambridge Research Systems, 2018). While participants completed the tasks, their head was stabilised with a SR research chin and forehead rest placed approximately 50cm from the monitor.

#### EEG experiment

Participants’ electrophysiological activity was recorded using a 64-channel BioSemi Active-Two system, with a sampling rate of 512Hz. Recordings were grounded using common mode sense and driven right leg circuit, electrodes were attached to a standard 64-electrode Biosemi EEG cap, with electrodes placed according to the international 10-20 system (Jasper, 1958). An additional eight external electrodes were affixed to participants’ skin: one on each mastoid, one above and below each eye, and one on the outer canthi of each eye. During recording, all electrode impedances were kept within +/− 50 μV.

#### Overall procedure

Participants completed the behavioural and EEG components on separate days. During both sessions, participants completed the task in a dimly lit room, while their head was placed upon a chinrest. The behavioural component took 2-2.5 hours to complete, and the EEG component took 10 minutes to complete (excluding EEG setup).

#### Illusion procedure

In a single session, participants were tested on eight MPIs. This involved the participants completing eight experimental blocks in random order, with a separate block for each illusion (Figure 1). The eight illusions tested were: the flash-lag effect (FLE; Nijhawan, 1994), the luminance flash-lag effect (LUM-FLE; Sheth et al., 2000), the Fröhlich effect (Fröhlich, 1924), the flash-drag effect (FD; Whitney & Cavanagh, 2000), the flash-grab effect (FG; Cavanagh & Anstis, 2013), the motion-induced position shift (MIPS; De Valois & De Valois, 1991), the twinkle-goes effect (TG; Nakayama & Holcombe, 2021), and the flash-jump effect (FJ; Cai & Schlag, 2001). The illusion procedure was identical to that used in Cottier et al. (2023) and as such, the illusion specific dimensions and procedures are not discussed here. The only change made compared to Cottier et al. (2023) is that 16 practice trials were added to the beginning of the Fröhlich effect. Prior to being assessed for each illusion, participants completed a Qualtrics survey which checked their understanding of the experiment instructions, and then completed practice trials until they demonstrated sufficient understanding of each illusion (e.g., in the FLE, if the flash was 20 degrees of polar angle in front of the moving target, we made sure that the participants were reporting the flash as ahead). The understanding of participants was checked after each practice trial. Participants were asked to maintain fixation upon a fixation point (subtending approximately 0.3 to 0.5 degrees of visual angle) in the centre of a grey background. Breaks with no time limit were provided after each experiment block, and halfway during each block. The experimental code will be made available upon publication at: https://osf.io/nc9mx/?view_only=db3992fb03b54b8086c94657b7e4b7c1.

### Resting state EEG

The resting state session was organised into 10 60-second trials, 5 trials for each condition (eyes-open and eyes-closed), sequentially alternating between conditions. All participants completed the trials in alternating order starting with an eyes-open trial. Participants were instructed to stay still and relaxed throughout the recording, keeping their chin on the chinrest. During the eyes-open trial, participants were told to fixate upon a white fixation dot in the centre of a grey background (RGB value = 128) and minimise blinking. During the eyes-closed trial, participants were told to keep their eyes closed until they heard a beep signalling the start of the next trial. At the end of each trial, participants could take as long as they needed before pressing ‘space’ to proceed to the next trial. The start of each trial was indicated by an auditory beep played through the computer speakers. Resting state data collection was conducted by several researchers and could occur before or after participating in a separate EEG study. Four participants that completed Cottier et al. (2023) were brought back to complete just the resting state EEG component. The remaining participants provided resting state data while also participating in other EEG studies.

### Analysis

#### Behavioural pre-processing

All data cleaning and analysis was conducted with MATLAB (v.R2023b; The MathWorks Inc., 2023). The analysis code will be made available upon publication at: https://osf.io/nc9mx/?view_only=db3992fb03b54b8086c94657b7e4b7c1. All behavioural data was cleaned and analysed using the analysis procedures outlined in Cottier et al. (2023). In brief, for each participant in each block the magnitude of the associated illusion was estimated. For the five illusions that used 1-up-1-down adaptive staircases (FLE, LUM-FLE, FE, FD, and TG), the illusion magnitude was calculated as the average difference between the points of subjective equality (PSE) for each direction of motion of the inducer or target (e.g, in the FLE (clockwise – counterclockwise)/2). Calculating the average difference ensures the illusion magnitude is not twice its true size. The PSE for each direction was calculated by averaging across all the staircases for that direction (e.g., leftwards vs rightwards motion). For each staircase, a PSE was calculated by averaging the final 20 trials for the FLE, LUM-FLE, and FD, and the final 10 trials for FE and TG due to fewer available trials.

The MIPS, FG, and FJ did not use adaptive staircases, and for these illusions the magnitude was represented as the mean difference between the reported position and the physical position, within each direction of motion. In illusions with staircases, participants were excluded if their staircases did not converge. The criteria for whether a staircase converged are discussed in each illusion-specific subsection below. For participants that participated in Cottier et al. (2023) and completed two sessions, their illusion magnitude was calculated as the average of each magnitude across sessions.

#### Flash-lag effect (FLE)

The FLE magnitude was calculated as the arc length distance in degrees of visual angle between the end of the target rod and the flash. This was done within each direction of motion (clockwise and counterclockwise), then averaged across motion directions. For this illusion, we considered staircases as not converged if the difference between the two staircases for a given motion direction (one initialized ahead and one initialized behind) was greater than 3.18 degrees of visual angle (15 degrees of polar angle). Six participants that completed a single session had staircases that failed to converge, and one participant that completed two sessions had staircases that did not converge. These participants were excluded. Of participants that completed two sessions, the staircases of three did not converge in one session, but did converge in the other. As such, their effect was calculated using the session where the staircases converged. Of the 61 participants that completed this illusion (18 completed two sessions), 7 participants were excluded from further analysis due to these staircase criteria. The final sample comprised 54 participants, 37 of which completed a single session of illusions.

#### Luminance flash-lag effect (LUM-FLE)

The LUM-FLE magnitude was calculated as the difference between the PSE of the luminance of the target circle and flashed circle, at the moment of flash onset. Staircases were considered not converged if within any luminance change direction, the difference between the staircases with opposite initial values was greater than 30% luminance contrast. Applying this criterion, 8 participants that completed a single session, and 1 participant that completed two sessions, were excluded from the analysis for this illusion. Three participants that completed two sessions of this illusion had staircases that did not converge in the first session but did converge in the second session. As a result, their LUM-FLE was calculated using the data from the second session. Additionally, one participant was excluded due to a data saving error. Overall, of the 61 participants that completed this illusion (18 completed two sessions), ten participants were excluded. The final sample size comprised 51 participants, 34 of which completed a single session of illusions.

#### Fröhlich effect

The Fröhlich effect was the arc length difference in degrees of visual angle between the physical starting position of the rod’s trailing edge and the vertical meridian. Consistent with Cottier et al. (2023), participants were excluded if they pressed the same key for at least 80% of the trials in two or more staircases, or if their staircases did not converge. Staircases were considered to have not converged if, within a single motion direction, the difference between staircases with opposite starting values remained greater than 8.25 dva (45 degrees of polar angle). Applying these exclusion criteria led to two participants that completed two sessions being excluded for having staircases that did not converge in either session. The final sample size comprised 59 participants, of which 16 participants had completed two sessions and 43 participants had only completed a single session of illusions.

#### Flash-drag effect (FD)

On each trial, the FD was calculated as the vertical distance in degrees of visual angle between the PSE of the target rectangles and the central fixation point. The effect for each participant was calculated as half of the average difference between the PSE each direction (PSE for grating moving downwards – PSE for grating moving downwards/2). Staircases were considered not converged if the final staircase values within a direction of motion differed by more than 3.5 dva. No participants failed the staircase exclusion criteria, so there we no exclusions, meaning that the final sample size for this illusion comprised 61 participants, 43 that completed a single session of illusions.

#### Flash-grab effect (FG)

The FG magnitude was operationalised as the arc length distance in degrees of visual angle between the target’s position and the position reported by the participant. This was averaged across all trials within each reversal direction (clockwise and counterclockwise), then across reversal. Positive errors represent displacements in the direction of reversal motion. Participants were excluded if they failed more than 20% of the attention check trials, or made invalid responses for more than 10% of the total trials (18 trials). Invalid responses were mouse responses not on the annulus on trials when the target was presented. Four participants that completed a single session were excluded for failing the attention check. One participant that completed two sessions was excluded for making too many invalid responses. Of the 61 participants that completed this illusion (18 completed two sessions), five participants were excluded. The final sample comprised 56 participants, of which 39 completed a single session of the illusions.

#### Motion-induced position shift (MIPS)

The illusory effect was represented as half of the average horizontal offset between upper and lower Gabors at the point that observers reported the two to be horizontally aligned. A trial was excluded as an outlier if the absolute magnitude of the effect was equal to or greater than 10 degrees of visual angle. Of those that completed only a single session, two participants had a single trial removed, and two participants had two trials removed. Of the 16 participants that completed two sessions, 7 participants had a single trial removed, and two participants had two trials removed. No participants were excluded from this illusion. However, due to technical issues accessing the laboratory, time constraints meant one participant was unable to complete this illusion. Therefore, the final sample size for this illusion comprised 60 participants, of which 42 participants had completed a single session of the illusions.

#### Twinkle-goes effect (TG)

The TG was operationalised as the difference between the PSE of the dynamic noise trials and the static noise trials. The PSE was calculated for each staircase averaged within direction, and averaged across directions. The effect reflected half of the mean horizontal offset from vertical alignment at the point of perceptual alignment. Staircases were considered not converged if within each direction of motion, staircases with opposite initial values had PSE differences greater than 1.48 DVA. This criterion resulted in excluding three participants who completed a single session and one participant who completed two sessions. One participant who completed two sessions had staircases that did not converge in their first session but had staircases that all converged in their second session. As such, only their session 2 data was used to calculate the effect. Overall, of the 61 participants, four were excluded, yielding 57 participants, 40 of whom completed a single session of the illusions, 18 that completed two sessions.

#### Flash-jump effect (FJ)

The FJ was operationalised as half the average difference between the height of the target bar and the reference bar at the instantaneous moment of the flash. Positive values indicated an illusory shift in the direction of the size change (i.e., a growing bar was perceived as taller than veridical). To reduce the influence of premature responses, trials were considered outliers and excluded from the calculation if the magnitude on that trial was more than 3 standard deviations different than that participant’s mean effect. Among those who completed a single session, application of this rule led to one trial being excluded for seven participants, and two trials being excluded for one participant. Four participants who completed two sessions had a single trial removed. Two participants who completed a single session failed all three attention checks and were excluded from further analysis. Overall, of the 61 participants (18 of whom had completed two sessions), excluding two participants yielded 59 participants, 41 of whom had completed only a single session of the illusions.

#### EEG pre-processing

EEG data was pre-processed using the EEGLAB toolbox (version 2024.0; Delorme & Makeig, 2004) in MATLAB (version R2023b; The MathWorks Inc., 2023). The raw data and channel spectra for the 19 parietal-occipital electrodes (P9, P7, P5, P3, P1, Pz, P2, P4, P6, P8, P10, PO7, PO3, POz, PO4, PO8, O1, Oz, O2) was manually inspected to identify and remove (and later interpolate, see below) channels that were flat-lined or excessively noisy, and unlikely to contain signal. The data was then re-referenced to the average signal of all the EEG electrodes, before being trimmed to contain only the parietal-occipital electrodes of interest. The data was down sampled to 256Hz, the baseline (dc offset) was removed, and then the data was bandpass filtered using a 1Hz high-pass filter and a 40Hz low-pass filter. The continuous EEG data was then split into ten distinct 62 second epochs, from 1 second before the start of the trial to 61 seconds after the start of the trial. This epoch length was chosen to mitigate the effect of edge artefacts on the data (Cohen, 2014). To clean the data, Independent Component Analysis (ICA) was conducted using the infomax algorithm implemented using the extended runica function in EEGLab. The ICLabel classifier was used to automatically label the ICA components, and automatically reject components that had a 90% or greater probability of being a muscle, eye, or heart artefact (Pion-Tonachini et al., 2019). Following ICA, the spherical spline method (Perrin et al., 1989) was used to interpolate removed channels. This resulted in a single channel being interpolated for eight participants, and two channels being interpolated for three participants. The epochs were then trimmed to only contain the 60 seconds from the beginning of the trial.

#### Calculating IAF

IAF was calculated using the automated method developed by Corcoran and colleagues (2018). This method applies an algorithm to get two measures of IAF: peak-alpha frequency and centre of gravity. The algorithm estimates the power spectral density using the MATLAB implementation (*pwelch.m*) of Welch’s modified periodogram method (Welch, 1967). Then, a Savitzky-Golay curve fitting method with a frame-width of 11, and a polynomial order of 5, was used within the alpha domain of 7-13Hz to smooth the power spectral density output before estimating the peak alpha frequency (PAF) and the centre of gravity (COG). The PAF is the frequency within the alpha band exhibiting the largest amplitude (Tarasi & Romei, 2024). The COG computes a weighted average of the power within the alpha band, representing the average activity of alpha oscillations (Goljahani et al., 2012). The COG is a good measure of IAF when there are multiple alpha peaks or no alpha peak present in the EEG spectra, making it difficult to compute a distinct PAF (Corcoran et al., 2018; Goljahani et al., 2012). Per the recommendations of Corcoran et al. (2018), in this study we report both measures of IAF.

For each measure of IAF and for each participant, we required an estimate of the measure of IAF for at least 9 channels, before averaging across channels. Using this criterion, for the eyes-open condition, there were 17 participants for whom PAF could not be estimated, and 12 participants whose COG could not be estimated. In the eyes-closed condition, all participants had at least 9 channel estimates for each measure, and PAF and COG could be estimated for all participants. Since reliable IAF estimates were not possible in the eyes-open condition, we restricted our analyses exclusively to the data from the eyes-closed condition. This is consistent with the fact that eyes-closed data is often preferred due to its greater test-retest reliability (Grandy et al., 2013). Within our eyes-closed condition, there was a strong significant positive correlation between PAF and COG (*r*_s_ = .95, *p* < .001), showing strong inter-measure reliability between the two IAF estimates.

### Statistical inference

The histograms for each illusion and measure of IAF indicated that the data was not normally distributed (Appendix Figure 1). This was confirmed by statistically significant Kolmogorov-Smirnov tests (Appendix Table 1) with p *<* .001. As such, consistent with our previous publication (Cottier et al., 2023), non-parametric statistical analyses were conducted, and 95% confidence intervals were calculated with bias-corrected and accelerated (BCa) bootstrapping (*N* = 1000; Efron & Tibshriani, 1994). Spearman’s Rho was used for the correlation analyses. Correlation estimates will always be attenuated by measurement noise (Mollon et al., 2017; Spearman, 1987). As such, to correct for this measurement error and get a “true” estimate of the correlations between IAF and the illusions, we calculated disattenuated correlations using Spearman (1987)’s formula (see also Cottier et al., 2023). Disattenuated correlations are reported alongside the regular “attenuated” correlations. However, we will not interpret the disattenuated correlations, as they are simply provided as an estimate of the true effect, and are not intended for inference (Hedge et al., 2018). To calculate the disattenuated correlations, we used the test-retest reliabilities for the illusions published by Cottier et al. (2023), and the test-retest reliability for Grandy et al. (2013)’s eyes-closed young IAF control group (.87).

**Table 1.** Mean and standard deviation for each illusion’s magnitude. The table shows data for the present study and for Cottier et al. (2023).

| Illusions | *M*(SD) |  |  |
| --- | --- | --- | --- |
|  | Unit of measurement | Present study | Cottier et al. (2023) |
| Flash-lag effect (FLE) | Degree of visual angle | 1.68 (2.14) | 1.70 (1.78) |
| Luminance flash-lag effect (LUM-FLE) | % Luminance contrast | 12 (12) | 12 (13) |
| Fröhlich effect (FE) | Degree of visual angle | 1.58 (1.53) | 1.2 (1.36) |
| Flash-drag effect (FD) | Degree of visual angle | 0.07 (0.07) | 0.06 (0.07) |
| Flash-grab effect (FG) | Degree of visual angle | 4.42 (1.51) | 4.34 (1.65) |
| Motion-induced position shift (MIPS) | Degree of visual angle | 0.84 (0.28) | 0.73 (0.25) |
| Twinkle-goes effect (TG) | Degree of visual angle | 0.34 (0.29) | 0.38 (0.26) |
| Flash-jump effect (FJ) | Degree of visual angle | 0.62 (0.48) | 0.44 (0.40) |

## Results

### Descriptive statistics

Figure 2 shows raincloud plots that provide the distribution, raw scores, range, median, and interquartile range for each illusion. These were created using Allen and colleagues (2019) MATLAB function. Overall, illusory effect magnitudes were qualitatively similar to the observations in Cottier et al. (2023; Table 1). Inspection of the raincloud plots (Figure 2) suggests that there might be individual differences present in the magnitude of each illusion, and in the measures of IAF. The mean PAF was 10.34 Hz (range = 8.58 to 12.21; *SD* = 0.83 Hz), and the mean COG was 10.16 Hz (range = 8.55 to 12.31; *SD* = 0.83 Hz). Inspection of the power spectra (Figure 3) confirms that a peak in the alpha band was present for all participants. PAF and COG were also strongly correlated (*r*_s_ = 0.95, *p* < .001, 95% BCa CI = [0.89, 0.98]).

### Illusion magnitudes and IAF estimates

**Figure 2.**
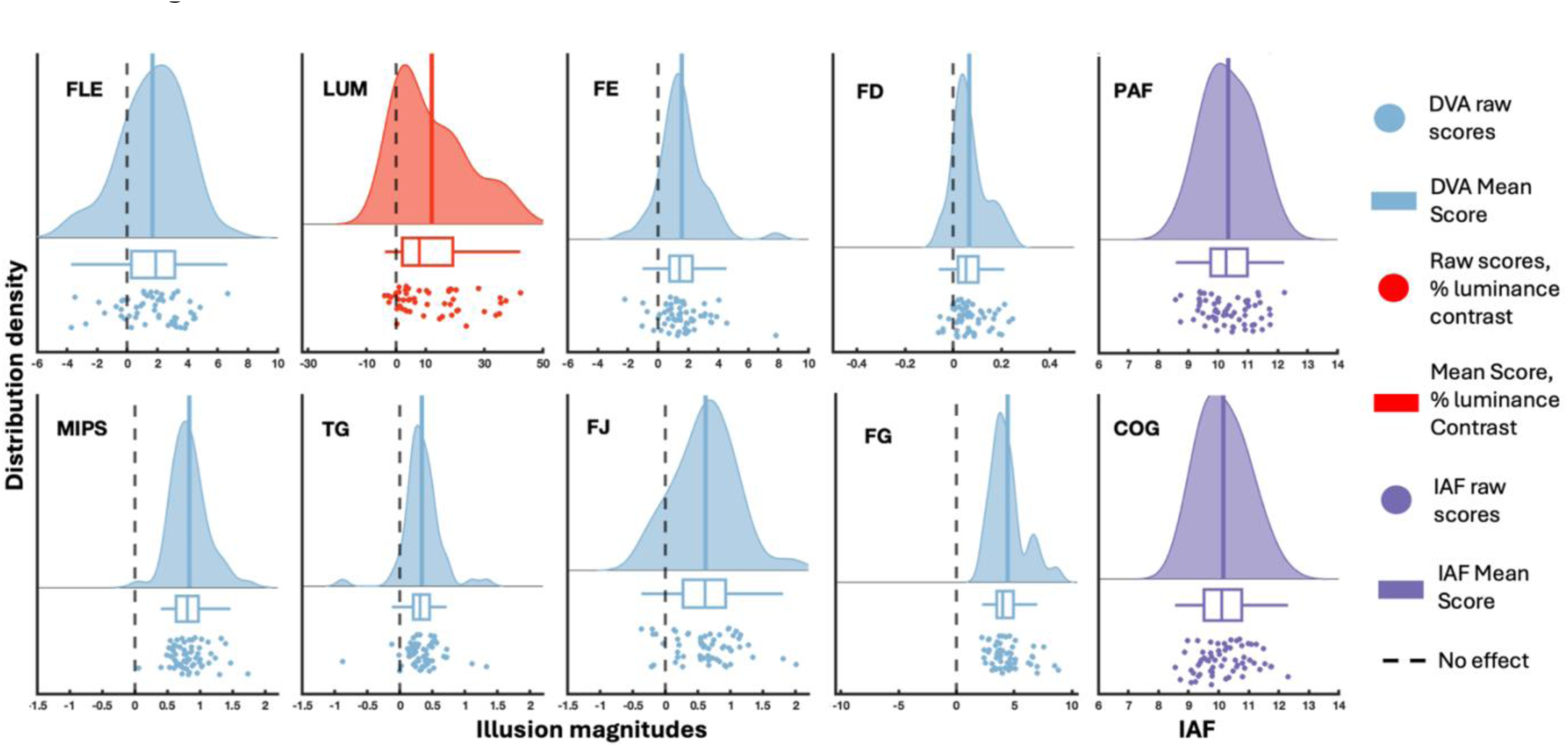
Raincloud plots for each illusion and the two IAF estimates. Blue colours show illusory magnitude measured in degrees of visual angle, red colours show illusory effect measured in % of luminance contrast, and purple colours show measures of IAF. The dashed black line shows the point corresponding to no illusory effect, with positive values representing an illusory effect in the expected direction. Boxplots show the interquartile range and median. The distributions show an estimated probability density distribution created using MATLAB’s ksdensity function with the mean marked with the solid vertical line.

### Correlation analyses

We calculated Spearman’s Rho correlation coefficients to explore whether individual differences in illusion magnitude were related to participants’ IAF estimates (Figure 4). Scatterplots of the relationship between illusions and the measures of IAF are presented in the Appendix (Figures 2 to 10). Biased corrected and accelerated (BCa) bootstrapped (*N* = 1000) confidence intervals are presented in Table 2. Bonferroni-Holm correction was used to control the family-wise error rate for multiple comparisons (Holm, 1979). The uncorrected p values for the correlation analyses are presented in Appendix Table 2. As shown in Figure 4, after Bonferroni-Holm correction we observed no statistically significant correlations between any of the illusions, or between any illusions and the measures of IAF. The only statistically significant correlation we observed was between COG and PAF. However, prior to Bonferroni-Holm correction statistically significant correlations were observed between the Fröhlich effect and flash-grab effect (*r*_s_ =.35, *p* = .009, 95% *BCa* CI = [0.07, 0.55]), and the twinkle-goes and motion-induced position shift (*r*_s_ = .34, *p =* .009, 95% *BCa* CI = [0.07, 0.54]). The former was not significant after correction for multiple comparisons in the data of Cottier et al. (2023), but the latter was.

**Figure 3.**
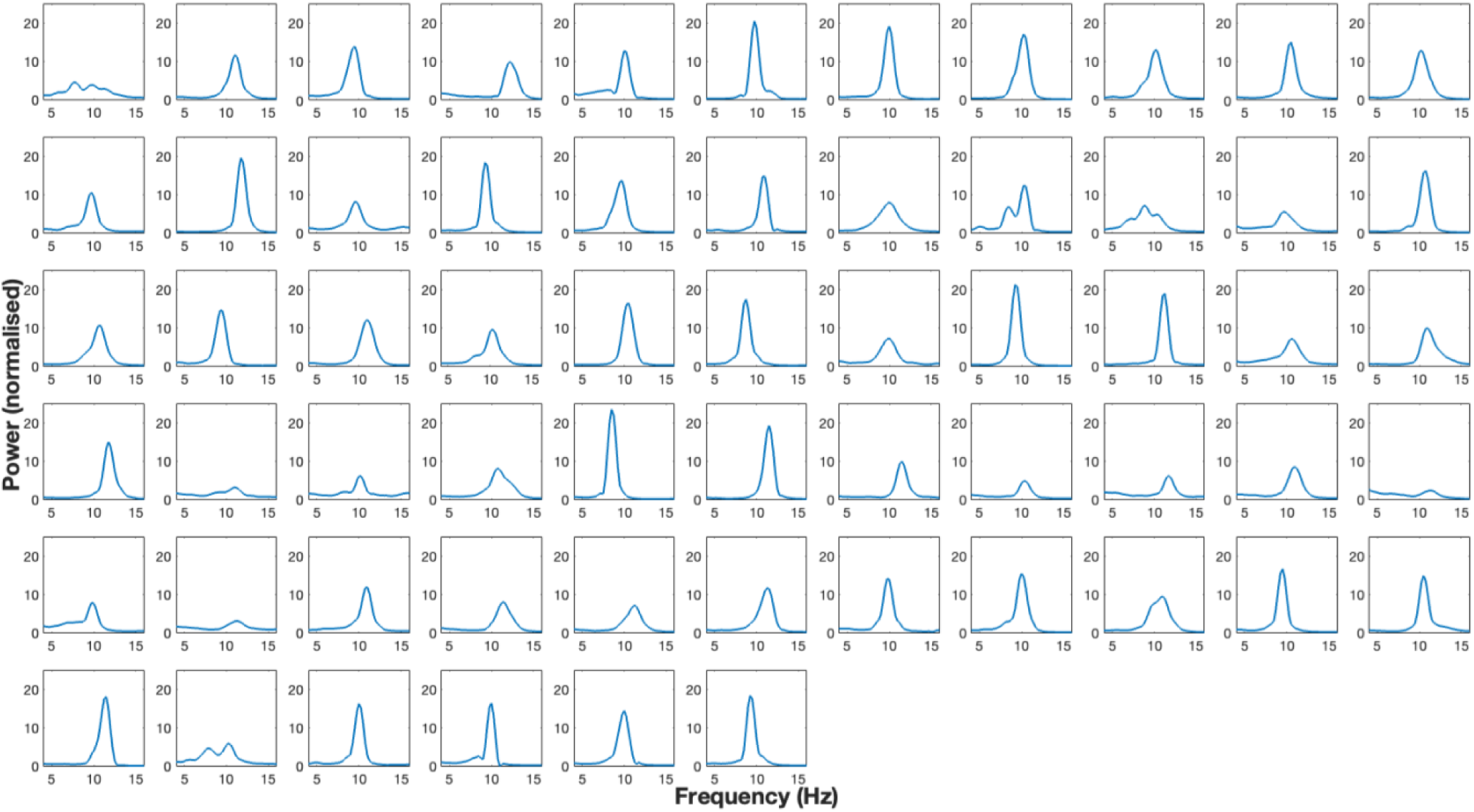
The Q-weighted power spectral density estimate for each participant. For each participant, the power spectrum was averaged across the power spectra for each channel. All participants experienced a peak in the alpha band (7-13hz).

**Figure 4.**
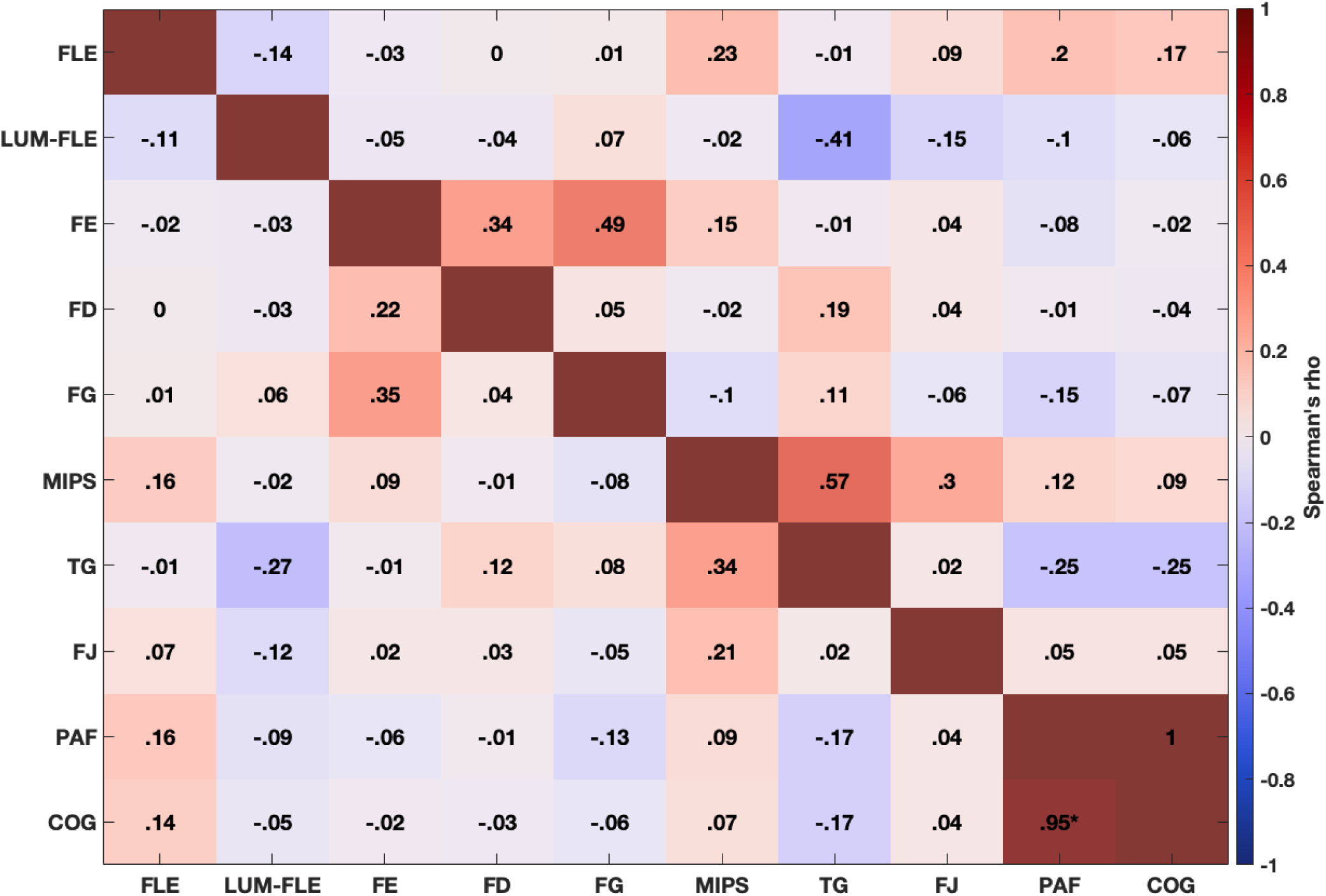
Correlation matrix showing the correlations between each illusion and the measures of IAF, PAF and COG. The disattenuated correlations are presented above the diagonal line, and the raw correlations are presented below the diagonal. The p-values for these correlations are presented in Appendix Table 2. Note, these correlations are not age controlled (for age-controlled correlations see Appendix Figure 11). Correlations using the data from a subset of occipital electrodes (O1, Oz, and O2) are provided in Appendix Figure 12. Statistically significant (*p* < .01) correlations are marked with an asterisk. The red diagonal boxes separate raw and disattenuated correlations.

**Table 2.** 95% bias-corrected and accelerated (BCa, *N* = 1000) bootstrapped confidence intervals for the correlations between illusions and IAF (IAF), using all parietal-occipital electrodes. The confidence intervals for the electrode subset O1, Oz, and O2 is presented in Appendix Table 3. Confidence intervals that do not contain zero are shown in **bold red font**. The blue cells show the confidence intervals for correlations not controlling for age. The red cells show the confidence intervals for correlations controlling for age.

| All parietal-occipital electrodes |  |  |  |  |  |  |  |  |  |  |
| --- | --- | --- | --- | --- | --- | --- | --- | --- | --- | --- |
| Illusions | FLE | Lum-FLE | Fröhlich | FD | FG | MIPS | TG | FJ | PAF | COG |
| FLE |  | [-0.4, 0.19] | [-0.34, 0.27] | [-0.35, 0.2] | [-0.3, 0.31] | [-0.1, 0.43] | [-0.29, 0.32] | [-0.17, 0.43] | [-0.04, 0.5] | [-0.07, 0.47] |
| Lum-FLE | [-0.4, 0.21] |  | [-0.31, 0.26] | [-0.31, 0.3] | [-0.22, 0.35] | [-0.31, 0.3] | **[-0.51, -0.005]** | [-0.37, 0.2] | [-0.4, 0.2] | [-0.36, 0.23] |
| Fröhlich | [-0.3, 0.27] | [-0.3, 0.25] |  | [-0.03, 0.46] | **[0.05, 0.57]** | [-0.15, 0.33] | [-0.25, 0.25] | [-0.23, 0.29] | [-0.3, 0.21] | [-0.26, 0.26] |
| FD | [-0.28, 0.27] | [-0.3, 0.27] | [-0.03, 0.45] |  | [-0.25, 0.24] | [-0.3, 0.31] | [-0.13, 0.46] | [-0.21, 0.36] | [-0.2, 0.33] | [-0.19, 0.28] |
| FG | [-0.27, 0.32] | [-0.19, 0.35] | **[0.07, 0.55]** | [-0.21, 0.28] |  | [-0.34, 0.18] | [-0.21, 0.35] | [-0.29, 0.23] | [-0.37, 0.18] | [-0.32, 0.25] |
| MIPS | [-0.13, 0.39] | [-0.32, 0.31] | [-0.14, 0.34] | [-0.28, 0.28] | [-0.32, 0.22] |  | **[0.07, 0.55]** | [-0.1, 0.46] | [-0.15, 0.31] | [-0.18, 0.31] |
| TG | [-0.26, 0.29] | **[-0.53, -0.02]** | [-0.27, 0.29] | [-0.18, 0.37] | [-0.22, 0.33] | **[0.07, 0.54]** |  | [-0.27, 0.28] | [-0.44, 0.05] | [-0.45, 0.07] |
| FJ | [-0.22, 0.38] | [-0.37, 0.17] | [-0.27, 0.28] | [-0.26, 0.32] | [-0.30, 0.20] | [-0.06, 0.48] | [-0.26, 0.26] |  | [-0.25, 0.28] | [-0.24, 0.27] |
| PAF | [-0.14, 0.46] | [-0.37, 0.24] | [-0.33, 0.21] | [-0.26, 0.24] | [-0.37, 0.15] | [-0.19, 0.33] | [-0.41, 0.1] | [-0.21, 0.3] |  | **[0.9, 0.98]** |
| COG | [-0.16, 0.45] | [-0.35, 0.25] | [-0.32, 0.24] | [-0.26, 0.2] | [-0.32, 0.2] | [-0.19, 0.30] | [-0.42, 0.09] | [-0.23, 0.29] | **[0.9, 0.98]** |  |

It has been noted that IAF varies with age, and is slower in older adults (Grandy et al., 2013). Thus, we wondered whether the absence of statistically significant correlations between the illusions and IAF was a consequence of not controlling for age effects with IAF. Therefore, we conducted a partial correlation to control for the effects of participants’ age (Appendix Figure 11; see Appendix Table 2 for *p*-values). The partial correlation replicated the corrected correlations above, with a statistically significant correlation between the two measures of IAF, but no significant correlations between any of the illusions or the illusions and IAF measures.

Previous studies that have found a correlation between visual perception and IAF often only analyse data from a specific subset of electrodes (e.g., O1, Oz, and O2; Cecere et al., 2015; Howard et al., 2017). In the present study, we analysed the data from 19 electrodes over the occipital and parietal cortex, making it possible that we could have been tapping into a mixture of oscillatory sources. Therefore, we repeated the correlation analysis using only the occipital electrodes typically used in in past research. Focusing the data on only three electrodes resulted in more missing data, as participants required an IAF estimate for all three channels of interest in-order to calculate the PAF and COG. As a result, PAF (*M* = 10.34, *SD* = 0.85, range = 8.5 to 12.32) could be estimated for 55 participants, and COG for 60 participants (*M* = 10.16, *SD* = 0.86, range = 8.23, 12.2). The non-age-corrected and age-corrected correlation matrices (Appendix Figure 12) replicated the patterns reported above, with the only significant correlation being between COG and PAF (see Appendix table 3 for confidence intervals). Overall, all four correlation analyses provide no evidence for a correlation between the two measures of IAF and any illusion magnitudes across eight different MPIs. This suggests that the magnitude of MPIs cannot be predicted using IAF.

### Correlations between illusions

Regarding the relationships between the illusions themselves, after Bonferroni-Holm correction, we did not replicate Cottier et al. (2023), who reported correlations between the Fröhlich and FD (.37), and between the TG, MIPS, and FG. Qualitatively, however, whilst not reaching significance after correcting for multiple comparisons, the pattern of correlations nevertheless appeared similar to those reported by Cottier et al (2023). For example, we observed a correlation coefficient of .34 between TG and MIPS. Comparatively, Cottier et al. (2023) observed a correlation of .39 between the TG and MIPS. Therefore, we were interested in exploring the extent to which the correlation estimates were similar across the two studies.

To explore the similarity in correlation estimates, we plotted the 95% bias-corrected and accelerated (BCa) confidence intervals for each study in Figure 5. These confidence intervals show that there is a great deal of similarity in the correlation estimates between the present study and Cottier et al. (2023). However, there are some deviations between studies. Notably, in the present study there is evidence for a correlation between the TG and LUM-FLE, and between the FG and Fröhlich effect. In Cottier et al. (2023), there was no evidence for these correlations. The correlation between the TG and LUM-FLE was not statistically significant, and the correlation between the FG and Fröhlich was not significant after Bonferroni-Holm correction. Overall, the correlation estimates seem to be quite consistent.

**Figure 5.**
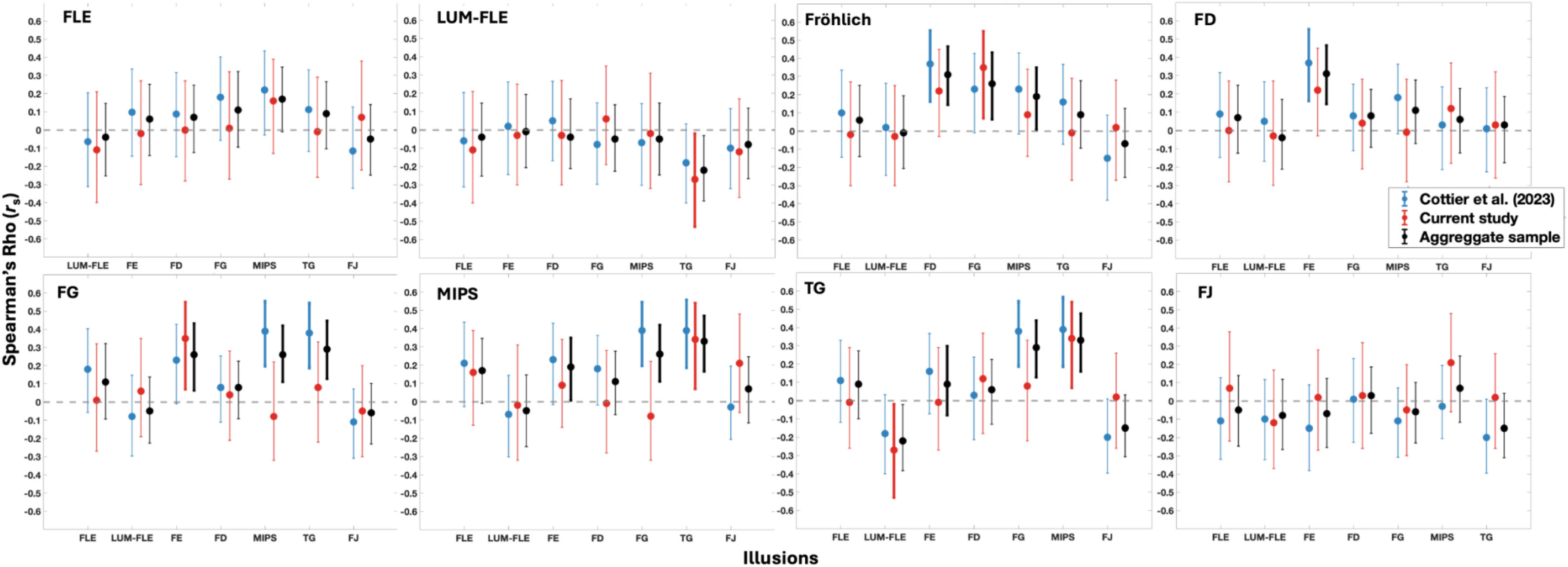
Error bars show the 95% Bias-corrected and accelerated (BCa) bootstrapped (*N* = 1000) confidence intervals for each illusion, for each study. Blue colours show the correlations and confidence intervals for Cottier et al. (2023). The red colours show correlations and confidence intervals for the present study. The black colours show the confidence intervals with the aggregate sample.

As we had new participants that completed the illusions from Cottier et al. (2023), we were interested in obtaining an updated version of the intercorrelation matrix presented by Cottier et al. (2023 – Figure 4). To this end, we created an aggregate dataset of 149 participants, comprising the 43 unique participants from the present study, and 106 participants from Cottier et al. (2023). The correlation analyses were repeated with this aggregate sample, and the updated correlation matrix is presented in Appendix Figure 13. The p-values for the updated correlation matrix are presented in Appendix Table 4. Overall, this auxiliary analysis replicated the key findings of Cottier et al. (2023). This is discussed in more detail in the Appendix materials. In Figure 5, we present the confidence intervals for the aggregate sample correlations. Ultimately, there is less variance in the confidence intervals for the aggregate correlations, indicating more precise correlation estimates. Overall, the conclusions to be drawn from this analysis are the same as those in Cottier et al. (2023) in that we observe evidence of weak to no correlation between the different illusions.

## Discussion

Examination of individual differences can allow us to better understand the mechanistic structure of visual perception. Previous work suggested a relationship between IAF and perceptual phenomena, leading to the suggestion that IAF may index the temporal resolution of perception. While some MPIs are thought to be related to temporal resolution (e.g., Linares et al., 2009), here we found no statistically significant correlations between IAF and eight different MPIs.

### Absence of correlations between IAF and motion-position illusions (MPIs)

The absence of correlations seems unlikely to be due to insufficient statistical power. Samaha and Romei (2024) found that the population correlation coefficient for the correlation between IAF and behavioural measures was typically between *r* = .39 to .53. Our sample size of 61 participants had 90% statistical power to detect relationships with a correlation coefficient above .37. Thus, our study was sufficiently powered to detect effects of the magnitude typically observed between IAF and behavioural measures (Samaha & Romei, 2024).

If a relationship between IAF and MPIs does exist, then its magnitude is likely to be much smaller than the relationship previously observed between IAF and other behavioural measures. Weak correlations could have been hidden by participants’ internal noise (Deodato & Melcher, 2024). For example, Deodato and Melcher (2024) found that they could only replicate the correlation between IAF and the two-flash fusion task reported by Samaha and Postle (2015) after using the slope of the psychometric function to control for participants’ internal noise. This suggests that participants’ internal noise can make it difficult to find a link between IAF and behavioural measures. The present study did not estimate participants psychometric functions, and is unable to implement this approach. As such, in the present study, it remains possible that participants’ internal noise may have masked weak correlations between IAF and MPIs. Future research could adopt Deodato and Melcher’s (2024) approach to minimise the effect noise may have on the correlation estimates.

Some of the tasks previously shown to correlate with IAF do not contain any motion, as they are cross-modal audio-visual tasks or tasks designed to estimate the thresholds of perception. Of those that do involve motion, possibly important differences remain (Howard et al., 2017; Minami & Amano, 2017; Ronconi et al., 2023; Shen et al., 2019; Zhang et al., 2019). For example, Ronconi et al. (2023) used the stream-bounce illusion, which is an audio-visual paradigm. The apparent motion Ternus display used by Shen et al. (2019) is a bistable stimulus. Zhang et al. (2019) used a bistable colour-motion feature binding task. It is possible that some aspect of these paradigms does correlate with IAF but is absent from MPIs. Additionally, in the case of Shen et al. (2019), they looked at pre-stimulus alpha before the task, whereas the present task looked at resting-state alpha, which may have a weaker correlation with behavioural tasks. Overall, it seems that although IAF is implicated in various aspects of visual perception, including motion tasks, it plays small to no role in MPIs.

### Absence of correlations between illusions

In our sample of 61 participants, after correcting for multiple comparisons we did not replicate the statistically significant correlations reported by Cottier et al. (2023). However, as shown in Figure 5, the correlation estimates were nevertheless highly similar across studies. A natural explanation for the absence of statistically significant correlations in the present study, is the smaller sample size in the present study (61 vs 106 in Cottier et al. (2023). However, statistically significant effects in Cottier et al. (2023) had correlation coefficients of 0.37 or higher and based on our sample size the current study had 90% power to detect effects of this size. However, participants completed fewer trials per illusion, viewing these illusions once, instead of twice as in Cottier et al. (2023), which increased the variability and effectively further reduced the statistical power. Therefore, it seems possible that the correlations between illusions might be truly smaller than reported in Cottier et al. (2023). This is supported by our confidence interval and correlation estimates, which show the estimated correlation with the aggregate sample was smaller than reported in Cottier et al. (2023).

### Discrete sampling is unlikely to account for MPIs

Based on the longstanding perceptual moment hypothesis (Stroud, 1967), Schneider (2018) proposed that discrete sampling could explain the FLE, Fröhlich effect, and other MPIs. Under the discrete sampling hypothesis for visual processing, the temporal resolution which IAF may index (Morrow & Samaha, 2022) would correspond to the duration of the visual system’s sampling window, and thus IAF should correlate with illusion magnitude (Morrow & Samaha, 2022). Our finding of no evidence for correlations between IAF and MPIs challenges the discrete sampling account of these illusions and suggests that this is not an underlying cause of these effects. This interpretation is corroborated by our observation that the FLE, Fröhlich effect, and the FJ did not correlate with one another, just as Cottier et al. (2023) and Morrow and Samaha (2022) found. Under discrete sampling, these illusions should correlate. Our results therefore suggest that discrete sampling at alpha is not involved in these illusions. However, we are not able to rule out the possibility that these illusions are driven by discrete sampling at different oscillation frequencies (Morrow & Samaha, 2022), or trial-level sampling processes which are independent from resting state mechanisms (see below). Furthermore, we cannot rule out the possibility of their being very small correlations between IAF and MPIs that this study was not sufficiently powered to detect.

Previous research has linked ongoing trial-level alpha dynamics (e.g., phase) to FLE magnitude (Chakravarthi & VanRullen, 2012; Chota & VanRullen, 2019). In the present study, we found no evidence for a link between trait-based components of alpha and the FLE. This difference in results may be due to the fact that the present study looked at resting state alpha dynamics recorded in a separate session to when participants completed the illusions, while previous studies have recorded EEG as participants complete the illusions. Thus, there could be some aspect of alpha (e.g., peristimulus phase) which is related to illusion magnitude, that the present study was not designed to detect. Given that peristimulus alpha dynamics (like phase) have been related to illusory perception (Cecere et al., 2015; Chakravarthi & VanRullen, 2012; Lange et al., 2014; Samaha & Postle, 2015) and that the position of moving objects can be decoded from ongoing trial-level alpha power (Turner et al., 2023), future research should explore how single-trial oscillatory dynamics mediate the perception of MPIs.

In conclusion, using an individual differences approach, the present study explored whether resting state individual alpha frequency (IAF) could predict the magnitude of eight motion-position illusions (MPIs). Correlation analyses found no evidence of an association between IAF and any of the illusions, suggesting that alpha-linked discrete sampling of visual information is not responsible for any of these effects. After correcting for multiple comparisons, we did not replicate the statistically significant effects reported in Cottier et al. (2023). However, bootstrapped confidence intervals revealed the correlation estimates were nevertheless highly similar across studies. An auxiliary analysis of aggregate data across these studies yielded updated, and more precise, estimates of inter-illusion correlations – overall showing evidence of weak to no association between these effects. Future research may explore how ongoing trial-to-trial oscillatory dynamics relate to MPIs. This would help to further characterise the extent to which neural oscillations influence motion and position perception.

## Data availability statement

Upon publication, the experiment code, analysis code, raw EEG data, and processed behavioural data will be made available at this link: https://osf.io/nc9mx/?view_only=db3992fb03b54b8086c94657b7e4b7c1.

## Author contributions

Timothy Cottier: Conceptualization, writing-original draft and review and editing, formal analysis, and investigation.

William Turner: Conceptualization, supervision, writing – review and editing.

Violet Chae: Investigation, writing – review and editing.

Alex Holcombe: Writing – Review and Editing.

Hinze Hogendoorn: Conceptualization, supervision, funding acquisition, writing – review and editing.

## Acknowledgements

The authors thank Luiza Bonfim Pacheco for assisting with resting state data collection.

## Appendix

**Appendix Figure 1.**
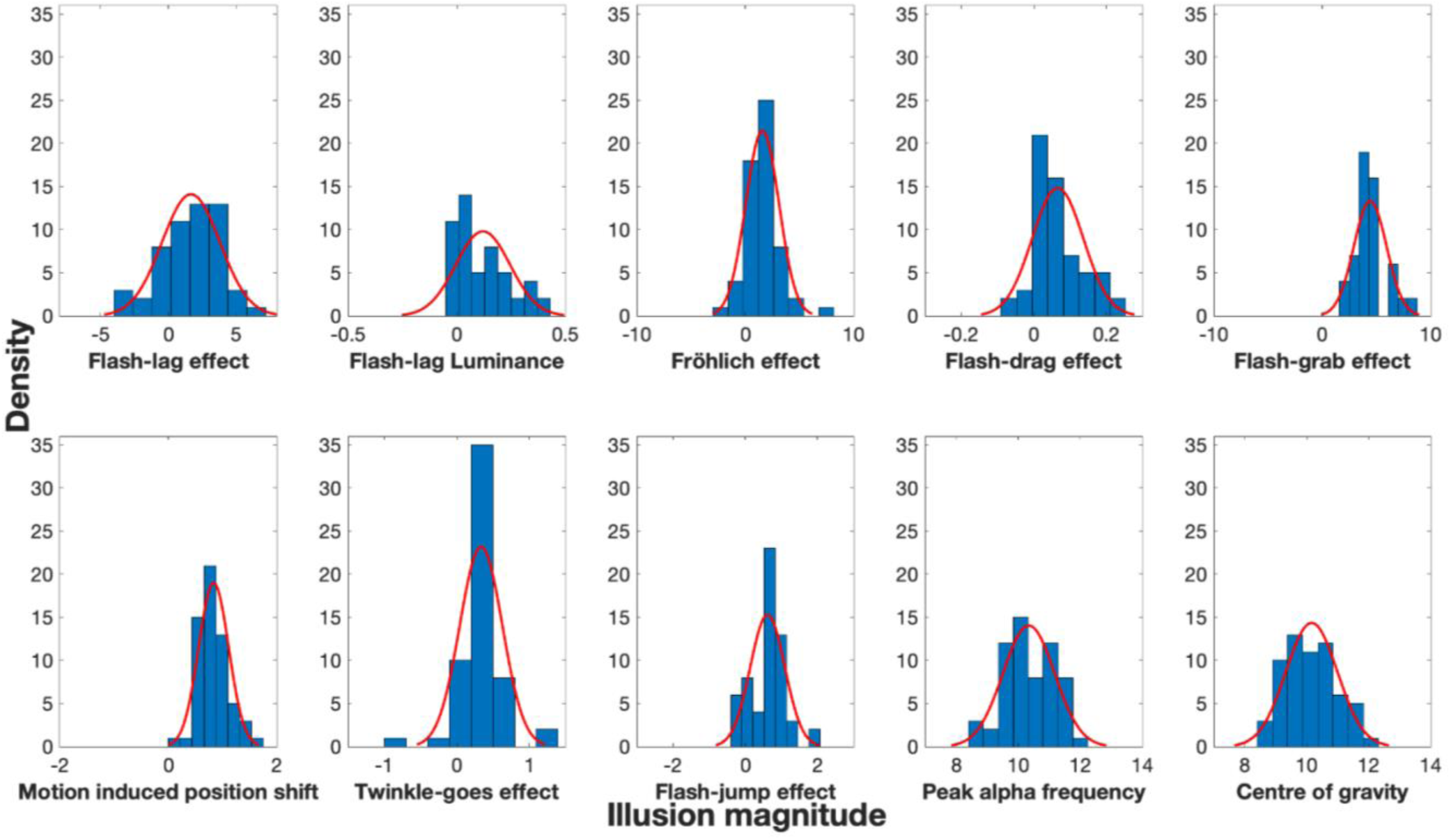
Histograms displaying the distribution for each illusion and IAF. Distributions were fit using the default parameters of MATLAB’s *“HistFit”* function.

**Appendix Table 1.** Kolmogorov-Smirnov tests for each illusion and individual alpha frequency (IAF). All Kolmogorov-Smirnov tests were significant (*p* < 0.001), suggesting the distributions were significantly different from a normal distribution.

| Illusions | Kolmogorov-Smirnov Statistic (*df*) |
| --- | --- |
| Flash-lag effect (FLE) | $D(53) = 0.52$ |
| Flash-lag luminance effect (LUM-FLE) | $D(50) = 0.48$ |
| Fröhlich effect (FE) | $D(58) = 0.55$ |
| Flash-drag effect (FD) | $D(60) = 0.48$ |
| Flash-grab effect (FG) | $D(55) = 0.99$ |
| Motion induced position shift (MIPS) | $D(59) = 0.66$ |
| Twinkle-goes (TG) | $D(56) = 0.46$ |
| Flash-jump (FJ) | $D(58) = 0.43$ |
| **Resting state EEG** |  |
| Peak Alpha Frequency (PAF) | $D(60) = 1$ |
| Centre of Gravity (COG) | $D(60) = 1$ |

**Appendix Figure 2.**
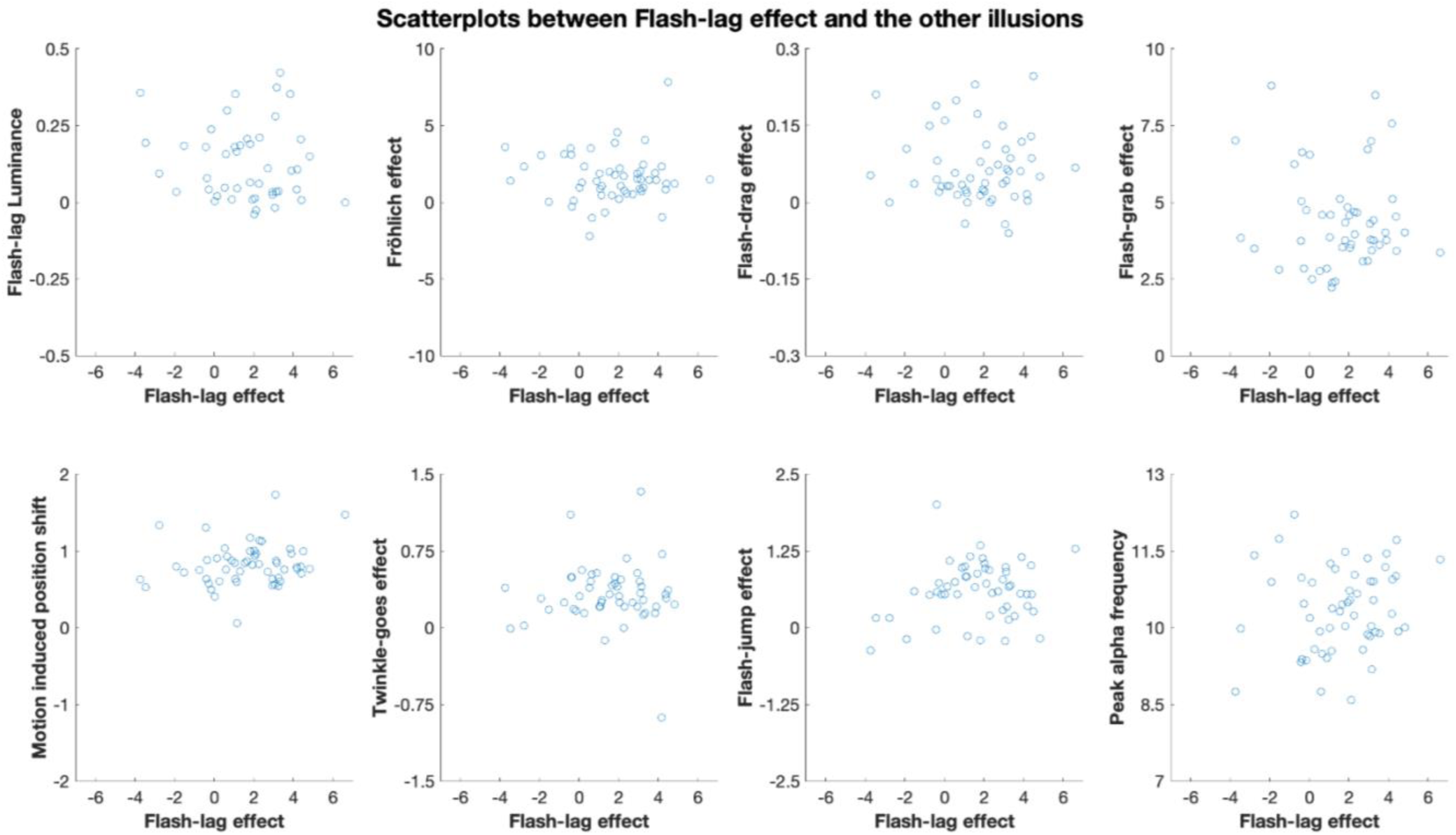
Scatterplots showing participants scores for the Flash lag effect, the other illusions, and peak alpha frequency.

**Appendix Figure 3.**
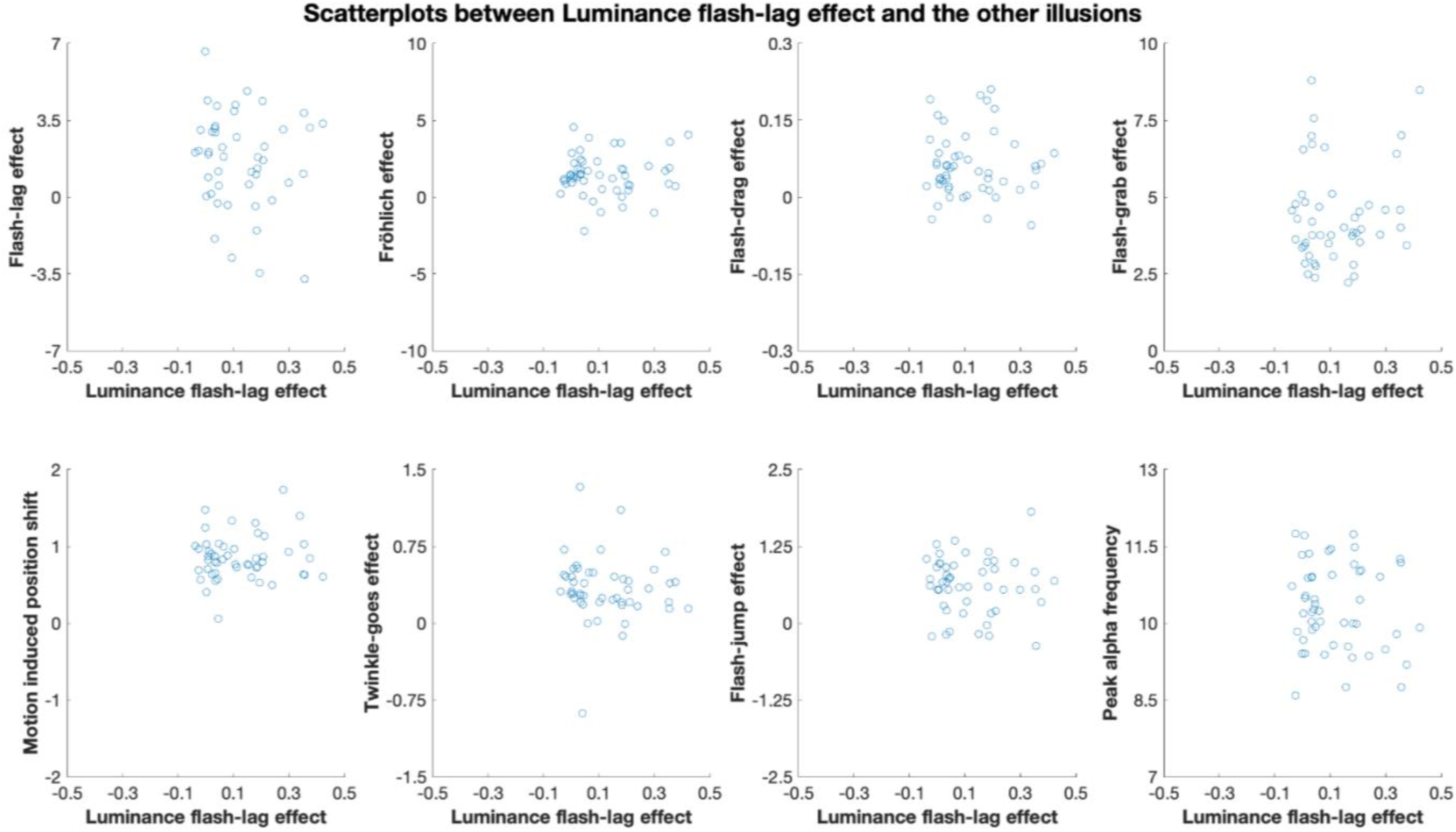
Scatterplots showing participants scores for the Luminance flash lag effect, the other illusions, and peak alpha frequency.

**Appendix Figure 4.**
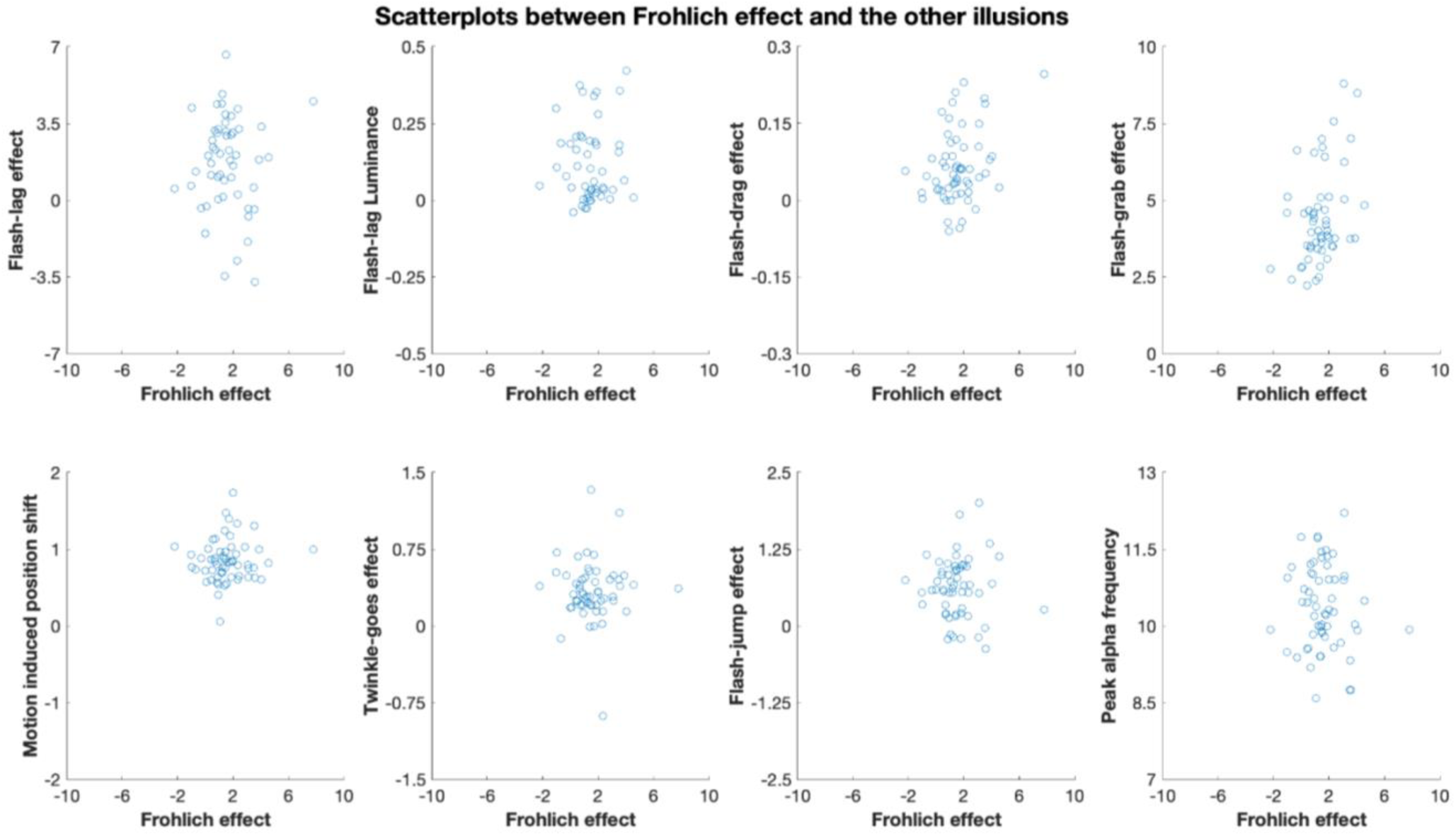
Scatterplots showing participants scores for the Fröhlich effect, the other illusions, and peak alpha frequency.

**Appendix Figure 5.**
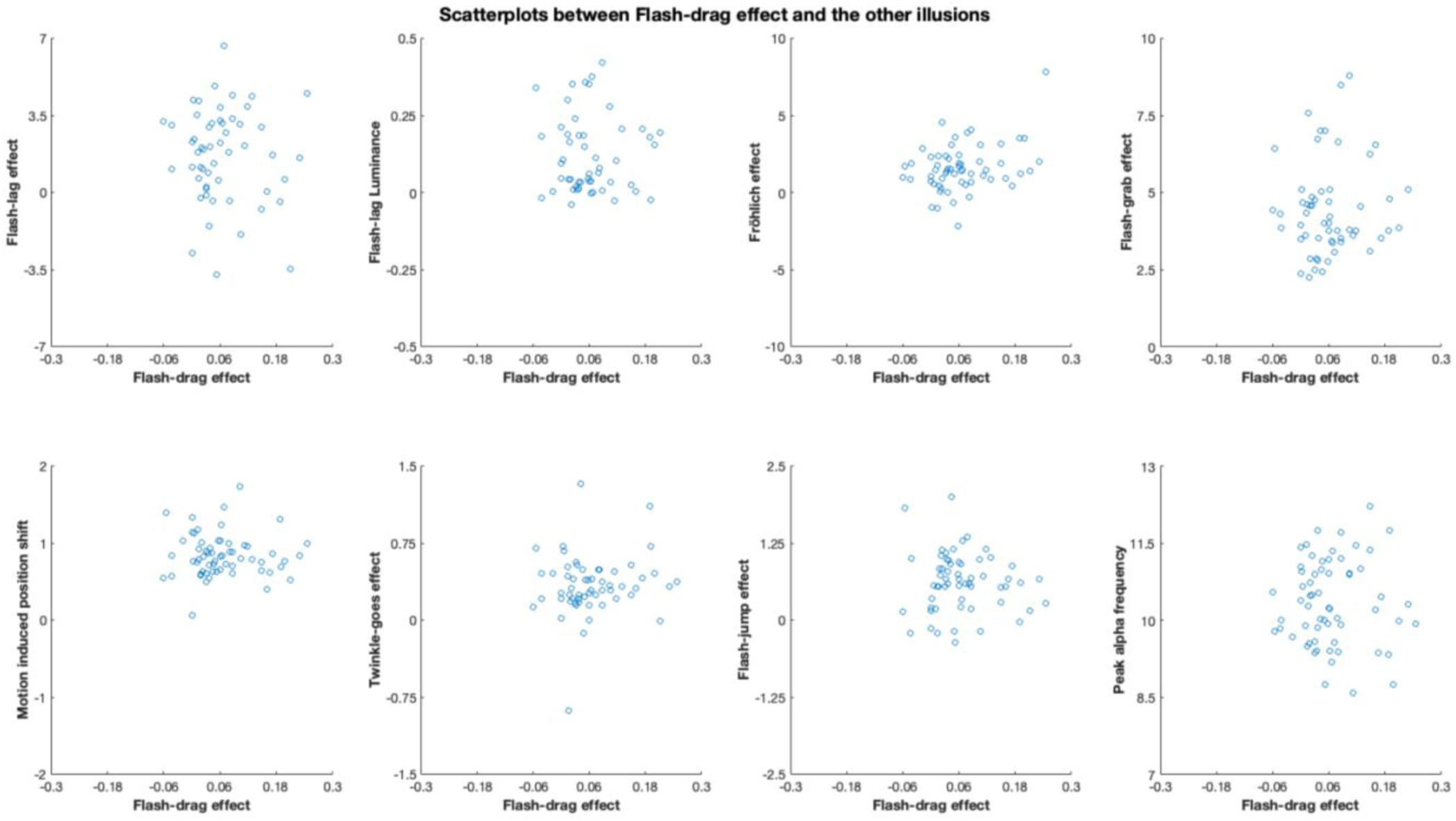
Scatterplots showing participants scores for the Flash-drag effect, the other illusions, and peak alpha frequency.

**Appendix Figure 6.**
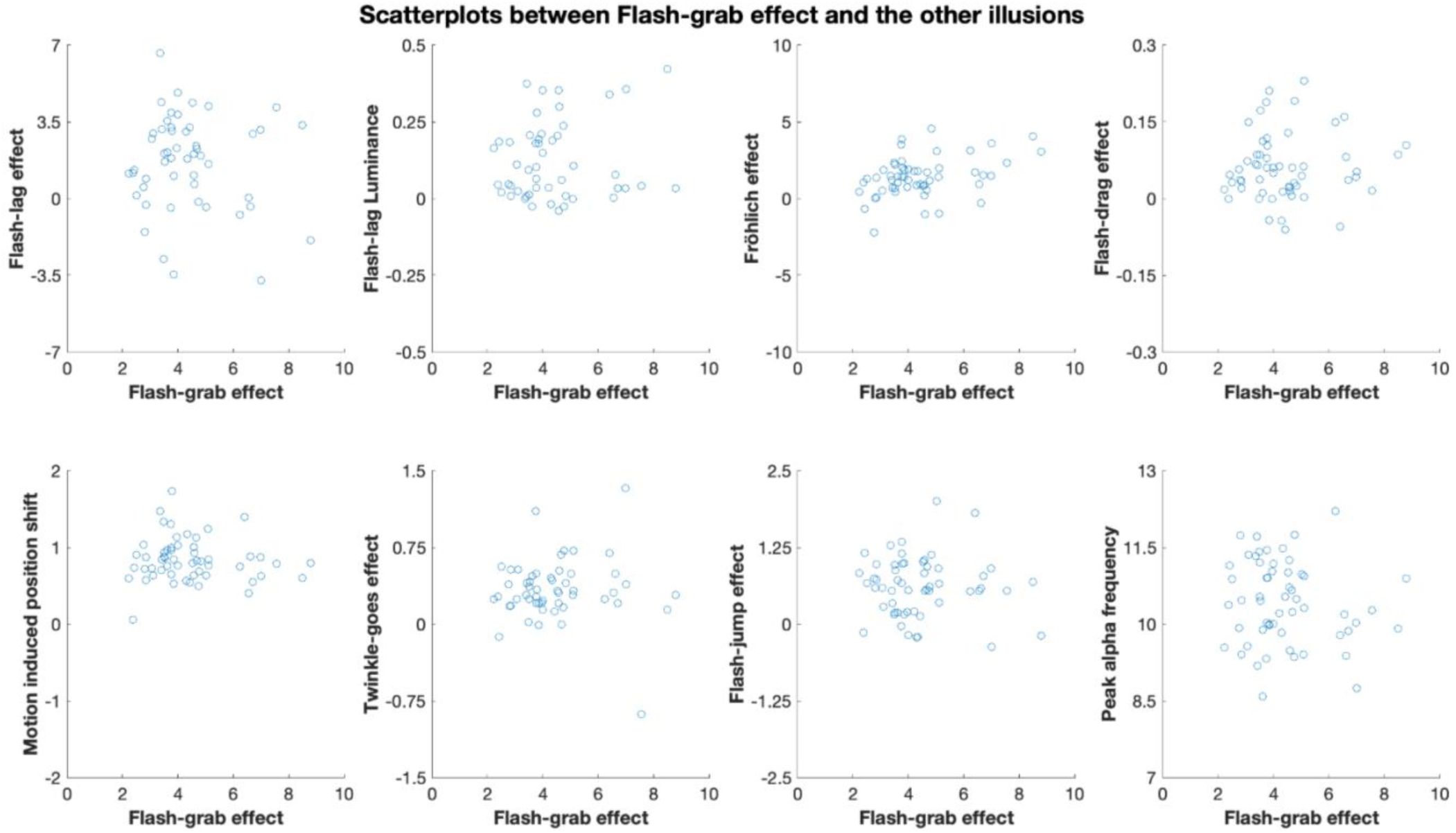
Scatterplots showing participants scores for the Flash-grab effect, the other illusions, and peak alpha frequency.

**Appendix Figure 7.**
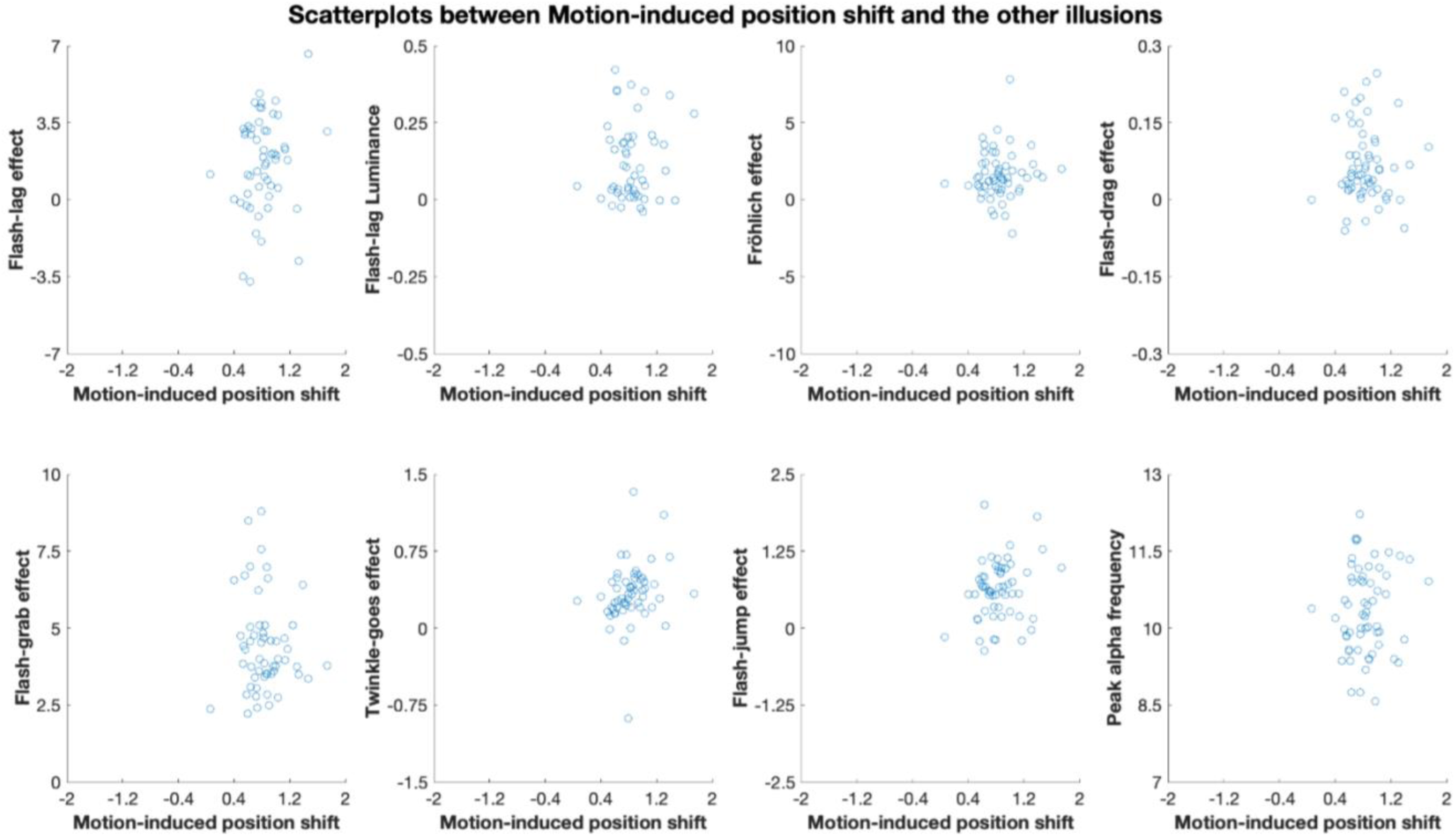
Scatterplots showing participants scores for the Motion-induced position shift, the other illusions, and peak alpha frequency.

**Appendix Figure 8.**
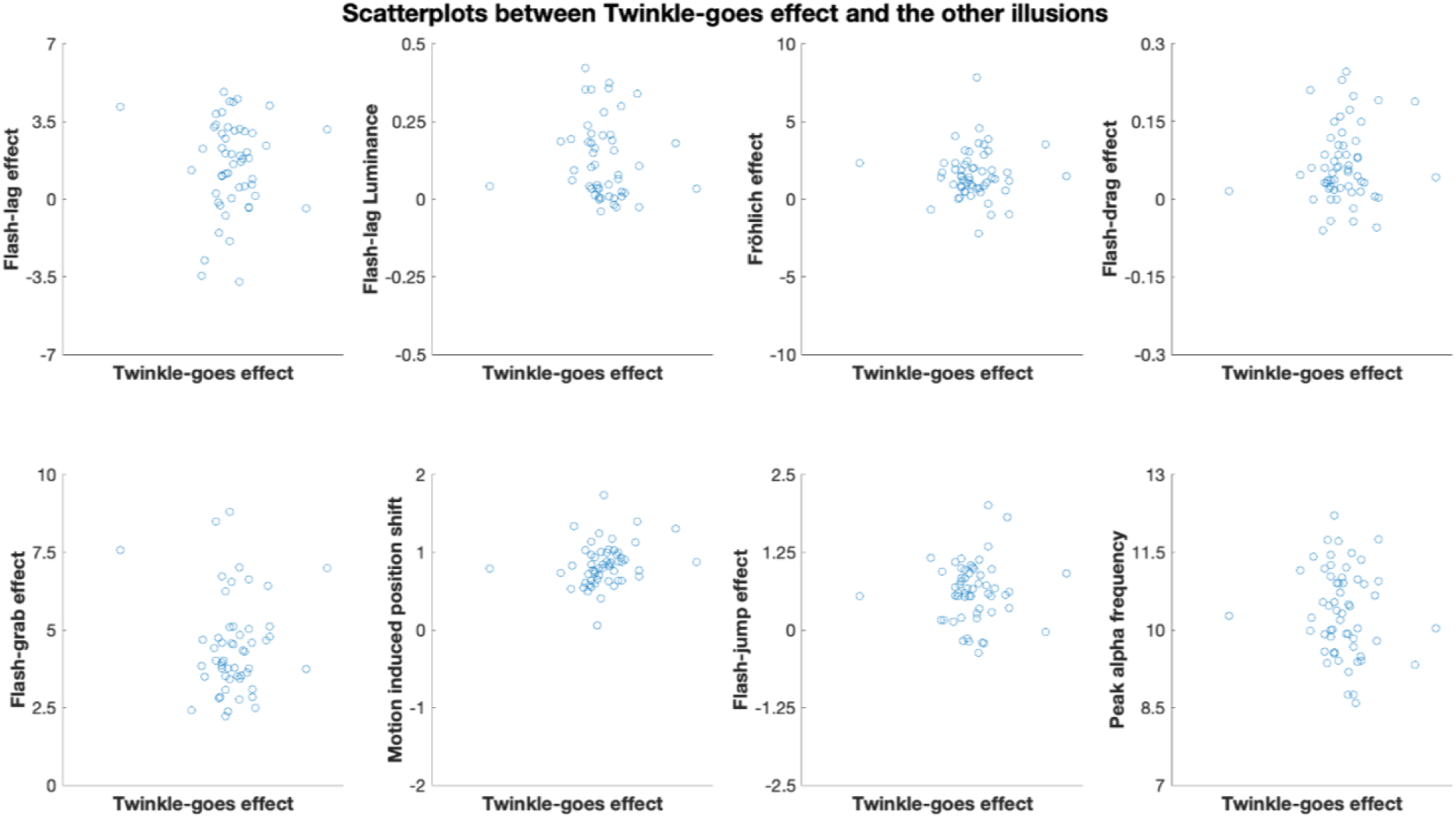
Scatterplots showing participants scores for the Twinkle-goes effect, the other illusion, and peak alpha frequency.

**Appendix Figure 9.**
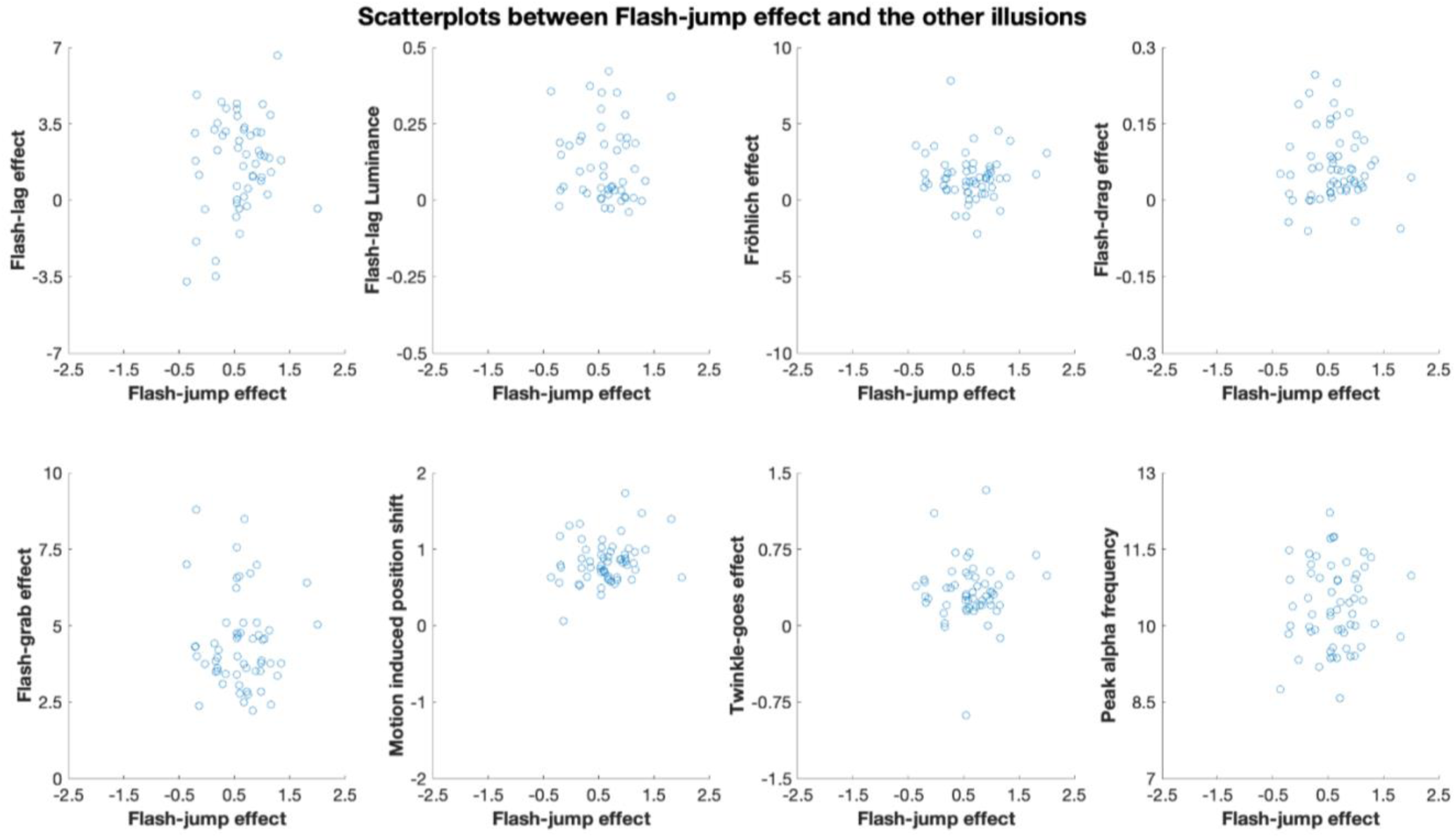
Scatterplots showing participants scores for the Flash-jump effect, the other illusions, and peak alpha frequency.

**Appendix Figure 10.**
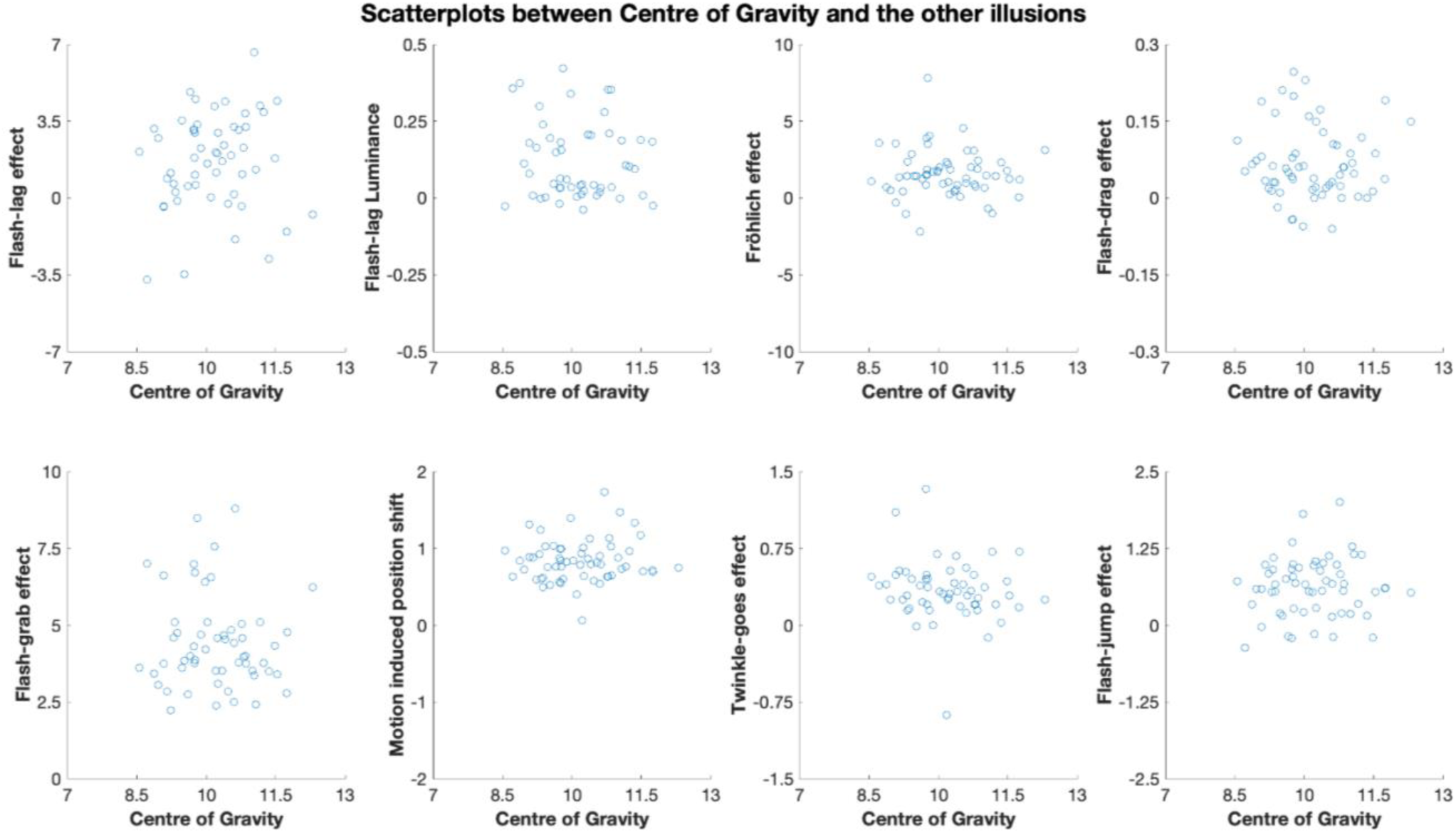
Scatterplots between participants Centre of Gravity and the other illusions.

**Appendix Table 2.**
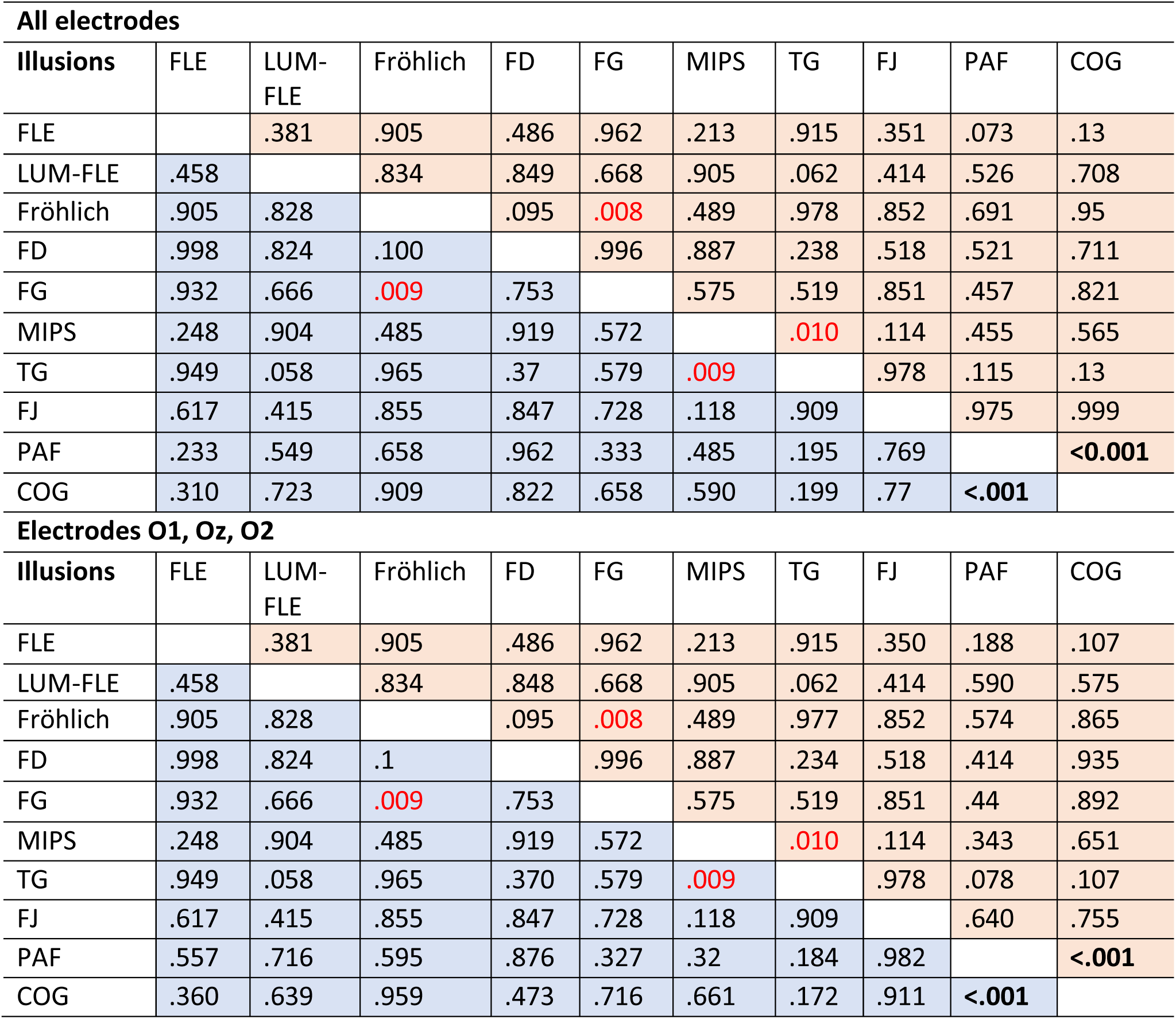
*P*-values for each correlation analysis between the illusions and IAF. The blue cells show the *p*-values for non-age corrected correlations, and the red cells show the p-values for age-corrected correlations. For the parietal-occipital electrodes, the non-age corrected correlations can be found in Figure 4, and the age corrected correlations are provided in appendix figure 11. The age corrected and non-age corrected correlations for electrode subset O1, Oz, and O2 are provided in Appendix Figure 12. Correlations statistically significant (*p* < .05) before Bonferroni-Holm correction, but not after Bonferroni correction are highlighted in red font. Correlations statistically significant after Bonferroni-Holm corrected are **bolded.**

**Appendix Figure 11.**
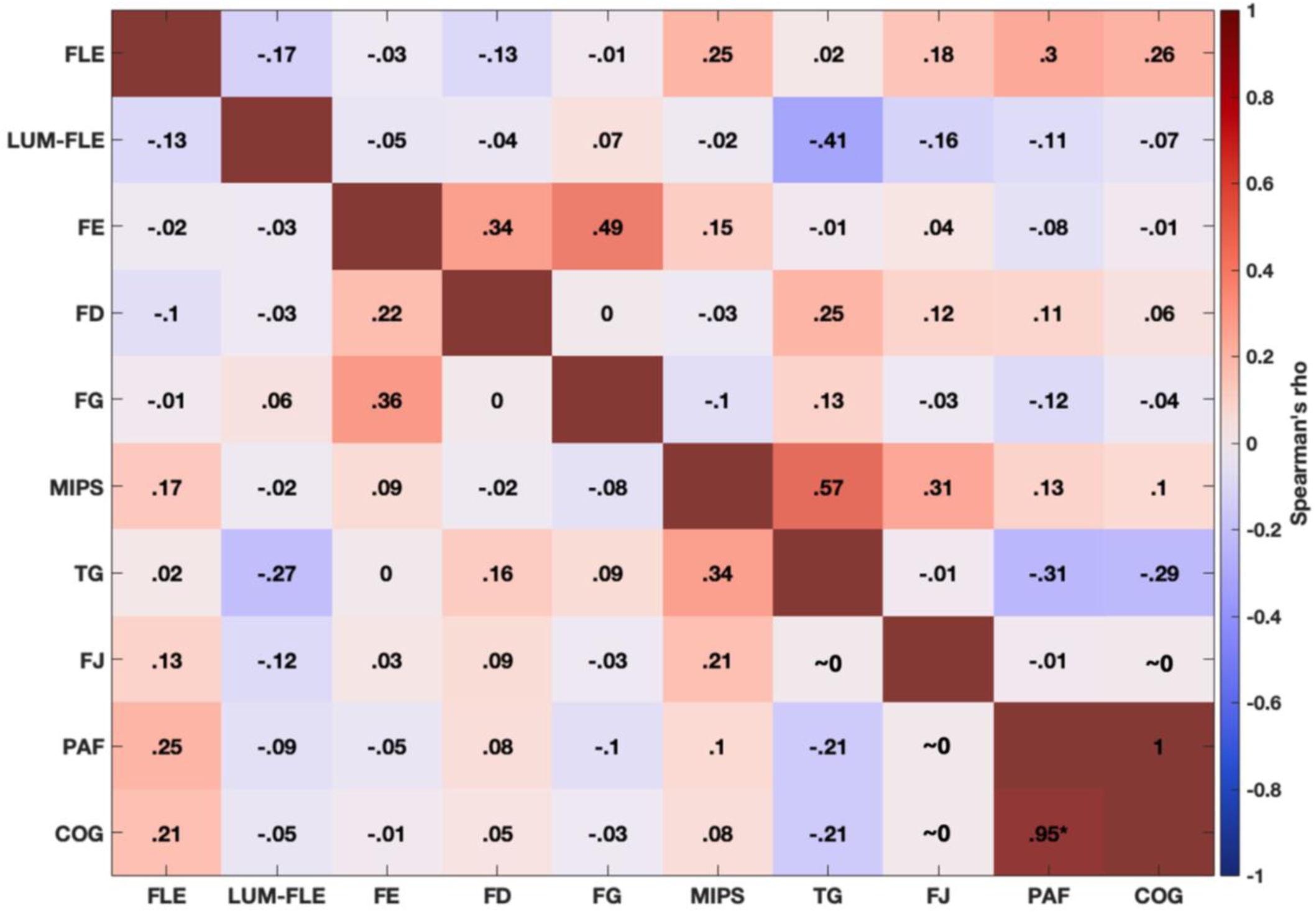
Correlation matrix showing the partial correlations controlling for age, between each illusion and the measures of IAF using all parietal-occipital electrodes. Correlations were rounded to two decimal places. FLE = Flash-lag effect, LUM-FLE = luminance flash-lag effect, FE = Fröhlich effect, FD = flash-drag effect, FG = flash-grab effect, MIPS = motion-induced position shift, TG = twinkle-goes effect, FJ = flash-jump effect, PAF = peak alpha frequency, and COG = centre of gravity. ∼0 = indicates that the correlation was less than < .01, and greater than >-.01 after the correlations were rounded to two decimals places. The disattenuated correlations are presented above the diagonal line, and the raw scores are presented below the diagonal. The p-values for these correlations are presented in Appendix table 2. Correlations not controlled for age correlations are presented in Figure 4.

**Appendix Figure 12.**
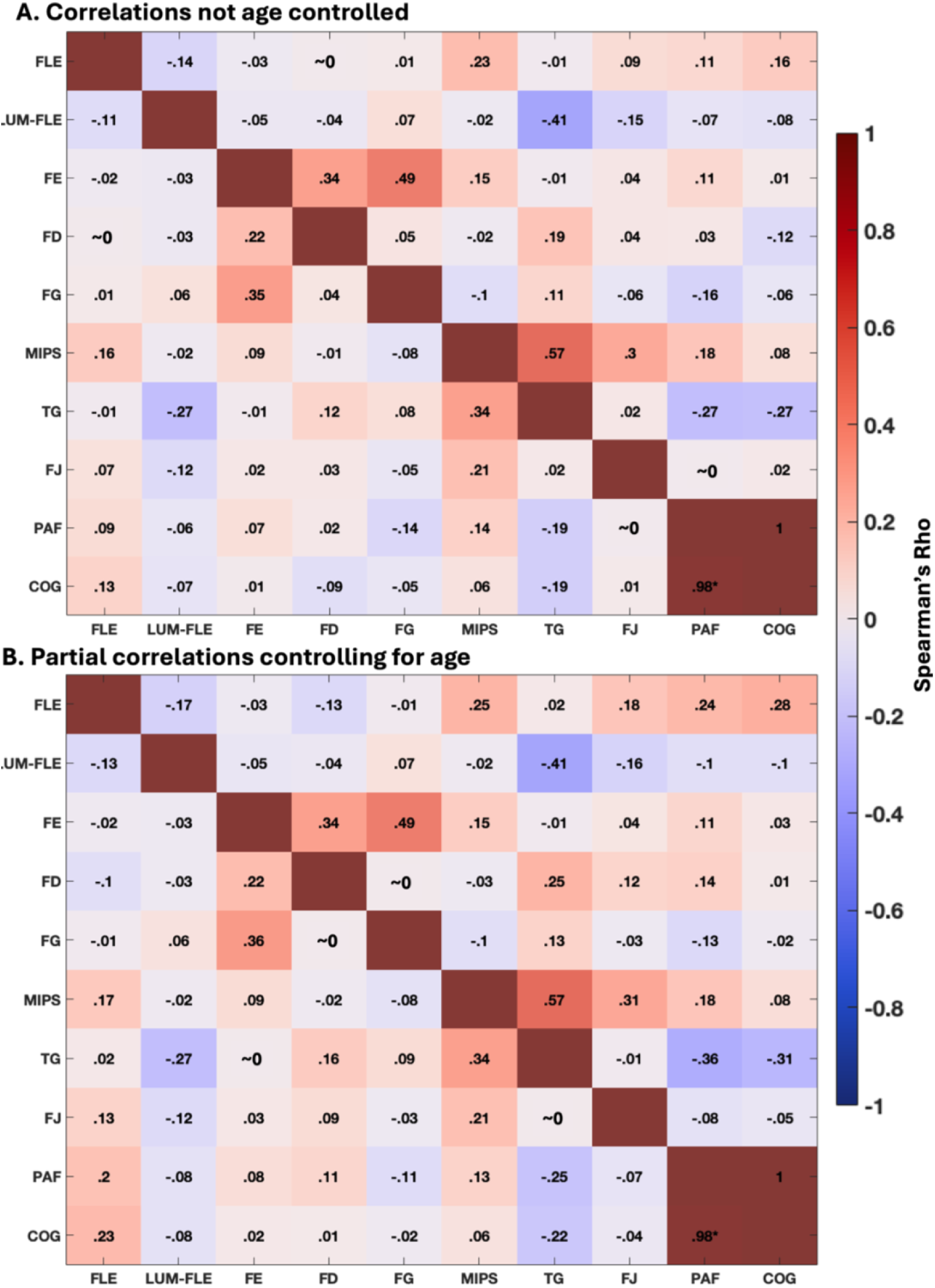
Correlations between IAF and the illusions, with IAF calculated with the data from a subset of electrodes (Oz, O1, and O2). Correlations were rounded to two decimal places. **A.** Show correlations before controlling for age. **B.** Shows correlations after controlling for age. FLE = Flash-lag effect, LUM-FLE = luminance flash-lag effect, FE = Fröhlich effect, FD = flash-drag effect, FG = flash-grab effect, MIPS = motion-induced position shift, TG = twinkle-goes effect, FJ = flash-jump effect, PAF = peak alpha frequency, and COG = centre of gravity. For the correlations between illusions including all parietal-occipital electrodes, see Figure 4 in main text. Disattenuated correlations are presented above the diagonal red line. ∼0 = indicates that the correlation was less than < .01, and greater than >-.01, after the correlations were rounded to two decimals places.

**Appendix Table 3.**
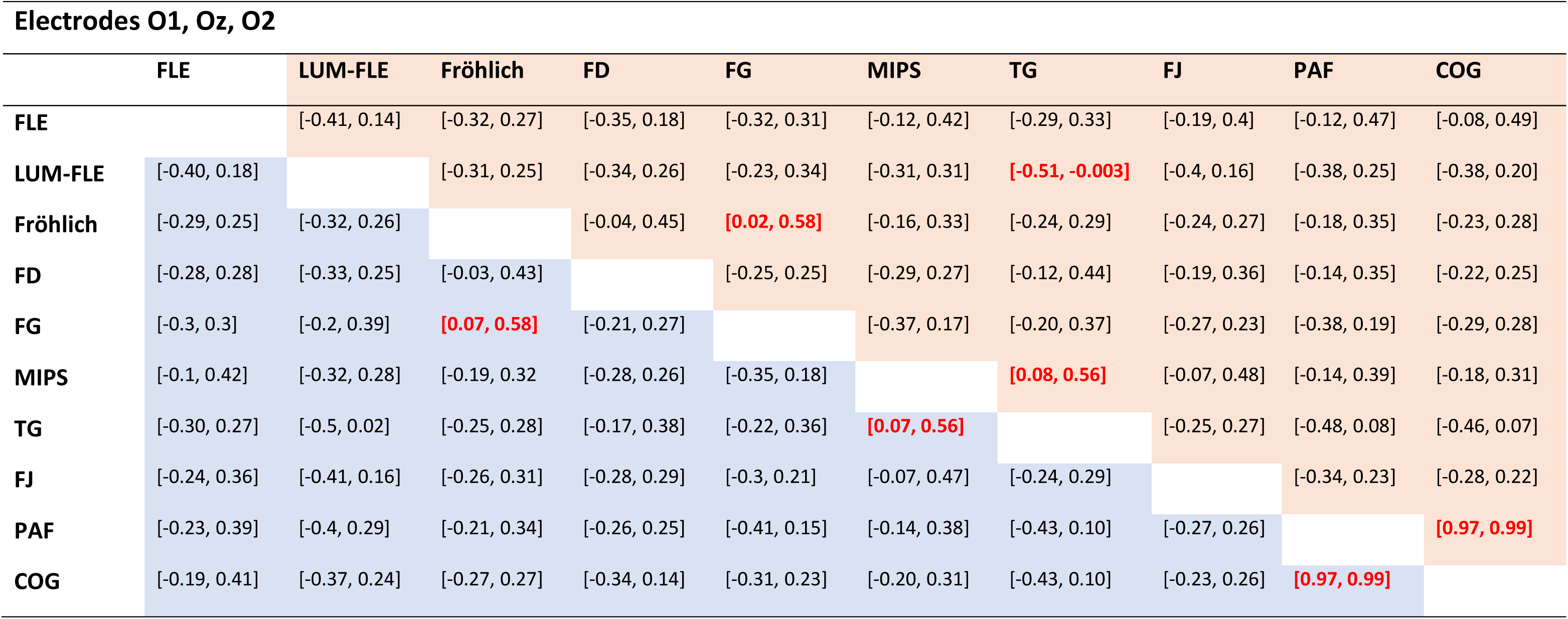
Bootstrapped confidence intervals for the correlations between illusions and IAF, when IAF is calculated only using the data from electrodes O1, Oz, and O2. Calculated using 95% bias-corrected and accelerated bootstrapping (*N* = 1000). Confidence intervals that do not contain zero are shown in **bold red font**. The blue cells show the confidence intervals for correlations not controlling for age. The red cells show the confidence intervals for correlations controlling for age.

|  | FLE | LUM-FLE | Fröhlich | FD | FG | MIPS | TG | FJ | PAF | COG |
| --- | --- | --- | --- | --- | --- | --- | --- | --- | --- | --- |
| FLE |  | [-0.41, 0.14] | [-0.32, 0.27] | [-0.35, 0.18] | [-0.32, 0.31] | [-0.12, 0.42] | [-0.29, 0.33] | [-0.19, 0.4] | [-0.12, 0.47] | [-0.08, 0.49] |
| LUM-FLE | [-0.40, 0.18] |  | [-0.31, 0.25] | [-0.34, 0.26] | [-0.23, 0.34] | [-0.31, 0.31] | **[-0.51, -0.003]** | [-0.4, 0.16] | [-0.38, 0.25] | [-0.38, 0.20] |
| Fröhlich | [-0.29, 0.25] | [-0.32, 0.26] |  | [-0.04, 0.45] | **[0.02, 0.58]** | [-0.16, 0.33] | [-0.24, 0.29] | [-0.24, 0.27] | [-0.18, 0.35] | [-0.23, 0.28] |
| FD | [-0.28, 0.28] | [-0.33, 0.25] | [-0.03, 0.43] |  | [-0.25, 0.25] | [-0.29, 0.27] | [-0.12, 0.44] | [-0.19, 0.36] | [-0.14, 0.35] | [-0.22, 0.25] |
| FG | [-0.3, 0.3] | [-0.2, 0.39] | **[0.07, 0.58]** | [-0.21, 0.27] |  | [-0.37, 0.17] | [-0.20, 0.37] | [-0.27, 0.23] | [-0.38, 0.19] | [-0.29, 0.28] |
| MIPS | [-0.1, 0.42] | [-0.32, 0.28] | [-0.19, 0.32] | [-0.28, 0.26] | [-0.35, 0.18] |  | **[0.08, 0.56]** | [-0.07, 0.48] | [-0.14, 0.39] | [-0.18, 0.31] |
| TG | [-0.30, 0.27] | [-0.5, 0.02] | [-0.25, 0.28] | [-0.17, 0.38] | [-0.22, 0.36] | **[0.07, 0.56]** |  | [-0.25, 0.27] | [-0.48, 0.08] | [-0.46, 0.07] |
| FJ | [-0.24, 0.36] | [-0.41, 0.16] | [-0.26, 0.31] | [-0.28, 0.29] | [-0.3, 0.21] | [-0.07, 0.47] | [-0.24, 0.29] |  | [-0.34, 0.23] | [-0.28, 0.22] |
| PAF | [-0.23, 0.39] | [-0.4, 0.29] | [-0.21, 0.34] | [-0.26, 0.25] | [-0.41, 0.15] | [-0.14, 0.38] | [-0.43, 0.10] | [-0.27, 0.26] |  | **[0.97, 0.99]** |
| COG | [-0.19, 0.41] | [-0.37, 0.24] | [-0.27, 0.27] | [-0.34, 0.14] | [-0.31, 0.23] | [-0.20, 0.31] | [-0.43, 0.10] | [-0.23, 0.26] | **[0.97, 0.99]** |  |

**Appendix Figure 13.**
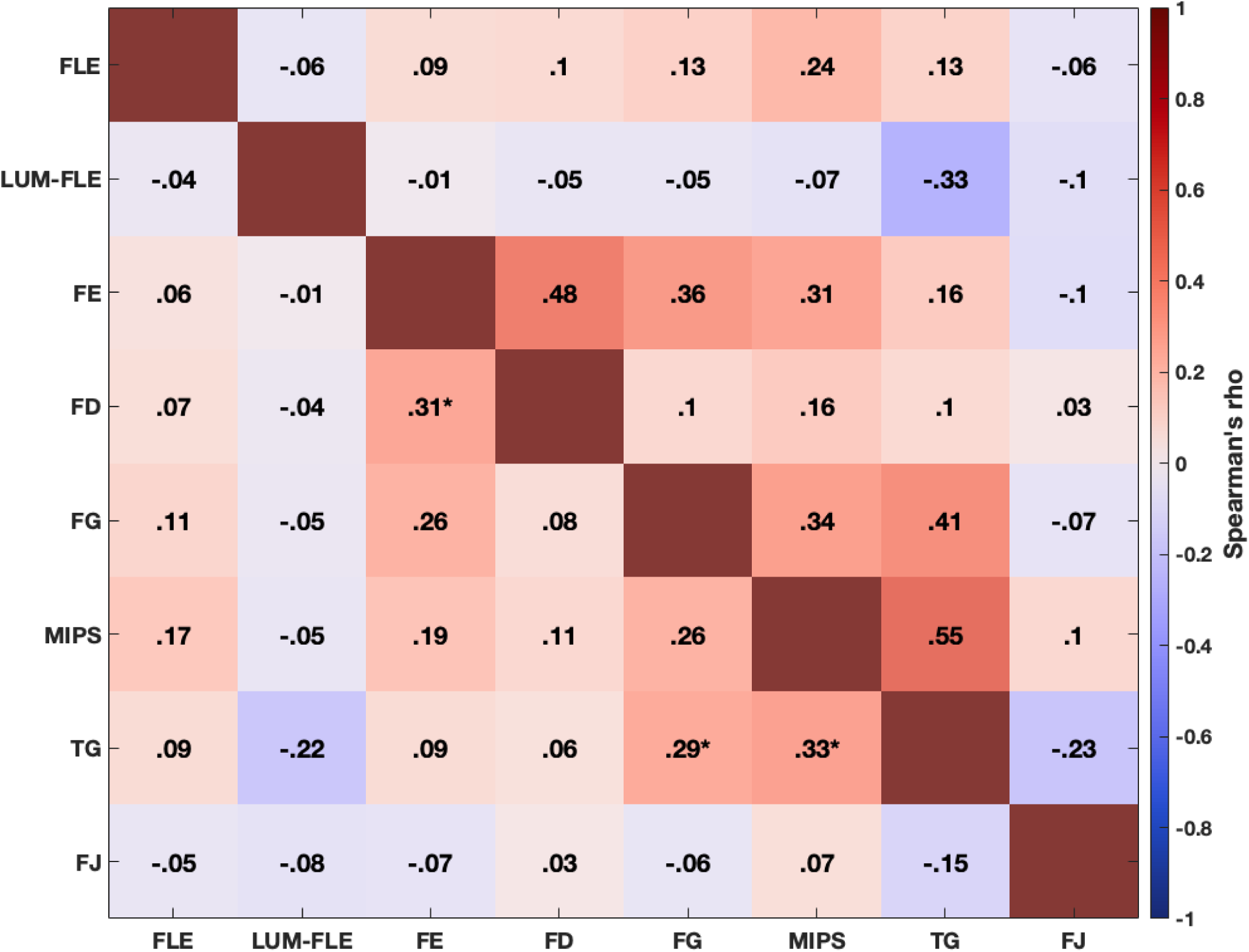
Correlations between the MPIs using the aggregate sample. The aggregate sample comprised 149 participants, 106 of which completed two sessions of the illusions in Cottier et al. (2023). We will refer to the participants from Cottier et al. (2023) as old participants. The sample size for each illusion in this correlation matrix comprised: 122 participants (85 old participants) completed the flash-lag effect (FLE), 117 (83 old) completed the luminance flash-lag effect (LUM-FLE), 125 (82 old) the Fröhlich effect (FE), 146 (103 old) completed the flash-drag effect (FD), 138 (99 old) completed the flash-grab effect (FG), 146 (104 old) completed the motion-induced position shift (MIPS), 136 (96 old) completed the twinkle-goes effect (TG), and 138 (97 old) completed the flash-jump effect (FJ). Disattenuated correlations are presented above the diagonal red line. The p-values for the correlations are presented below, in Appendix Table 4.

**Appendix Table 4.** P-values for the correlation analysis between the illusions, using the aggregate sample. The aggregate sample comprised the illusion magnitudes from the present study, and the illusion magnitude averaged across sessions from Cottier et al. (2023). Statistically significant p-values after Bonferroni-Holm correction for multiple comparisons are in **bold.**

| Illusions (n) | FLE | LUM-FLE | Fröhlich | FD | FG | MIPS | TG |
| --- | --- | --- | --- | --- | --- | --- | --- |
| FLE (122) |  |  |  |  |  |  |  |
| LUM-FLE (117) | .6674 |  |  |  |  |  |  |
| Fröhlich (125) | .5527 | .9477 |  |  |  |  |  |
| FD (146) | .4172 | .6953 | **.0005** |  |  |  |  |
| FG (138) | .2352 | .6349 | .0038 | .3447 |  |  |  |
| MIPS (146) | .0661 | .5915 | .0325 | .1980 | .0021 |  |  |
| TG (136) | .3543 | .0198 | .3218 | .4706 | **.0008** | **.0001** |  |
| FJ (138) | .6131 | .4267 | .4623 | .7720 | .5300 | .4114 | .0966 |

### Discussion of the aggregate sample correlations

As mentioned in the main text, 149 participants were included in the aggregate sample. 43 unique participants from the present study, and 106 from Cottier et al. (2023). Because participants in Cottier et al. (2023) completed the illusions twice across two separate sessions, we averaged their illusory effects across sessions. One of the critiques of Cottier et al. (2023), is that the conservative Bonferroni correction used to control for multiple comparisons might not have detected some true correlations. To address this possibility, we controlled for multiple comparisons by conducting the less conservative Bonferroni-Holm correction.

The correlation analysis with this aggregate sample replicated all but one of the observations of Cottier et al. (2023). Consistent with Cottier et al. (2023), we observed the same two correlated clusters of illusions. One cluster comprising the FD and Fröhlich effect, and another cluster comprising the TG, MIPS, and the FG. The only difference to Cottier et al. (2023) was that the correlation between FG and MIPS did not reach significance (p=0.0021; corrected alpha 0.002). Overall, we were not able to replicate the findings of Cottier et al. (2023) with the present study’s participants, which suggests that the effect might be smaller than originally reported.

